# The transcriptional regulator SutA is part of a nutrient scavenging network expressed at the entry to stationary phase in *Pseudomonas aeruginosa*

**DOI:** 10.64898/2026.06.26.734693

**Authors:** Claudia M Hemsley, Laurent Delavaine, Megan Bergkessel

## Abstract

Bacteria in natural environments frequently encounter nutrient limitation leading to growth arrest and must balance the potential benefits of continuing to respond to the environment by making new proteins against the costs of depleting limited resources. We previously showed that the RNA polymerase-binding regulator SutA enhances transcription of hundreds of genes during nutrient limitation in *Pseudomonas aeruginosa*, suggesting that it might be part of a regulatory network facilitating limited new protein synthesis. Here, we sought to expand our understanding of this network by identifying transcriptional regulators influencing *sutA* expression. Using northern blotting, western blotting, and reporter assays, we found that the sigma factors FliA and RpoS, and the DNA-binding regulator Lrp, impact expression from a proximal *sutA* promoter during the transition to stationary phase. This constellation of regulators and the dynamics of SutA expression lead us to propose that SutA is part of a regulatory network that facilitates scavenging. Scavenging includes motility toward possible nutrient sources and uptake mechanisms for these nutrients, activities which require an investment of resources but can yield important benefits during starvation. *In vitro* transcription experiments, proteomic analysis and reporter assays suggest that SutA directly supports new protein synthesis driven by RpoS and indirectly supports flagellar motility, perhaps by helping maintain protein biosynthetic capacity against the metabolic costs of motility. SutA expression is controlled by multiple regulatory inputs, including negative autoregulation, and the protein appears to be short-lived. These properties are consistent with a role in supporting short, controlled bursts of gene expression during nutrient limitation.

**Author Statement:** Many bacteria engage in cycles of colonising a nutrient-rich location, using the available nutrients, and then dispersing in search of a new location to colonise. While searching for new nutrients in a low-resource environment, bacteria will be starved and must coordinate resource-intensive processes such as new protein synthesis, motility, and nutrient uptake so that each crucial activity can be accomplished but none use too much of the limited pool of resources. We previously identified a regulator in *Pseudomonas aeruginosa* called SutA, which facilitates new protein synthesis under starvation conditions. Here, we have identified regulators of SutA expression. We find that the housekeeping sigma factor RpoD drives expression during growth, but at the entry to stationary phase, where SutA has obvious impacts on cellular physiology, the stress sigma factor RpoS, the flagellar sigma factor FliA, and the amino acid sensing transcription factor Lrp are important. Finally, we find that all cells in a nutrient-limited population express some SutA, but appear to do so in infrequent bursts, and that the protein is likely unstable. Together, these findings suggest that SutA contributes to the coordination of resource use while bacteria scavenge for new nutrients, facilitating limited amounts of new protein synthesis.

## Introduction

*Pseudomonas aeruginosa* is an opportunistic pathogen that can cause difficult-to-treat infections [1] and also colonise a wide range of environmental niches [2]. Its adaptability to diverse lifestyles may depend in part on its ability to tolerate and recover from resource limitation. *Pseudomonas* species have been observed to rise to dominance when nutrient-limited soils are supplemented, suggesting an ability both to persist when nutrients are low and to rapidly take advantage of new resources [3, 4]. In surveys of multiple bacterial species during starvation, *Pseudomonas spp.* have displayed a propensity to maintain both protein synthesis [5] and motility [6] at low levels where other organisms, including *Escherichia coli*, enter a more complete dormancy. *E. coli* has long been the primary model organism for studying responses to nutrient limitation by Gram-negative bacteria, so observations that *P. aeruginosa* respond differently suggest that explorations of regulatory mechanism during starvation in this organism could yield novel insights. Regulatory networks that contribute to maintenance of sustainable, low levels of activity during resource limitation in *P. aeruginosa* could contribute to its success in diverse niches, including infections.

To identify important regulators of starved states in *P. aeruginosa*, previous studies have explored the composition of the newly synthesised proteome and the genes contributing to fitness during prolonged total deprivation of external oxygen, carbon, or nitrogen sources [7–9]. Oxygen and carbon deprivation limit energy for powering cellular activities, while carbon and nitrogen deprivation limit the availability of biosynthetic building blocks to support growth. While these reductionist experiments deprived planktonic cells at moderate density of one resource at a time, *P. aeruginosa* likely experiences dynamic changes in resource availability during a natural cycle of colonising a niche, growing to high density, and then dispersing in search of a new high-resource niche [10]. Homologues of well-studied *E. coli* stress response regulators, such as the general stress sigma factor RpoS [11], the stringent response regulator DksA [12], and the heat-shock associated protease FtsH [13], appeared to be both robustly synthesised and important for fitness in one or more of these starvation conditions [7, 8]. However, unexpectedly, the flagellin protein FliC, present in thousands of copies in a flagellum, and many ribosomal proteins and translation factors continued to be among the most highly produced proteins during starvation [7]. This is perhaps consistent with the observed maintenance of protein synthesis and motility by starved *P. aeruginosa*, but at odds with the widely accepted *E. coli* paradigm that resource-intensive activities like ribosome and flagellar biogenesis should be strongly suppressed during starvation [11, 14]. Interestingly, mutations in *fleN*, encoding a negative regulator of flagellar biosynthesis in *P. aeruginosa*, caused strong fitness defects in carbon and nitrogen starvation [7], suggesting that tight regulation of ongoing investment in such activities is crucial.

Another protein that was among the most highly synthesised during oxygen and carbon deprivation was an uncharacterised protein subsequently named SutA for “<u>S</u>urvival <u>U</u>nder <u>T</u>ransitions”, after it was shown to be required for fitness during repeated transfers between anoxic growth arrest and rapid aerobic growth in lysogeny broth (LB) [9]. Deletion of the *sutA* gene also caused defects in biofilm formation, overproduction of the redox-active phenazine pyocyanin, and modest downregulation of hundreds of genes, including those encoding many of the ribosomal proteins, rRNAs, tRNAs, chaperones, core RNA polymerase (RNAP) subunits, and sigma factors. Chromatin immunoprecipitation (ChIP-Seq) experiments showed that SutA localised to many of these genes through its association with RNAP, suggesting that its role might be to support biosynthetic capacity by enhancing expression of the transcription and translation machinery during slow growth or starvation [9]. *In vitro* biochemical experiments showed that SutA is a small, mostly unstructured protein that binds directly to the β1 domain (or protrusion) of the RNA polymerase and enhances initiation at proximal rRNA promoters by both the stress sigma factor (RpoS) holoenzyme Eσ^S^ and the housekeeping sigma factor (RpoD) holoenzyme Eσ^70^, with a stronger effect observed for Eσ^S^ [15, 16]. Structural studies showed that SutA can influence the position of the β lobe domain of Eσ^S^ in complex with the rRNA promoter, favouring formation of the open complex [16]. Although Eσ^S^ has a distinct regulon that includes many genes not efficiently transcribed by Eσ^70^, the consensus sequence it recognises at the −10 promoter element is highly similar to that recognised by Eσ^70^, and it can initiate transcription from housekeeping promoters *in vitro*, albeit at lower levels [17]. Together, these observations suggest mechanisms by which SutA can support expression of housekeeping genes under starvation conditions when transcription may be challenged by changes to chromosome conformation, substrate availability, and the complement of available RNAP holoenzymes [15]. During protracted limitation for resources, such an enhancer of gene expression should be tightly controlled to prevent excessive investment of resources into biosynthetic activities that cannot be supported.

In this work, we have sought to identify regulators of SutA expression to gain a deeper understanding of the regulatory network that permits and controls ongoing investment in resource-intensive activities during nutrient deprivation in *P. aeruginosa*. Previously, a reporter construct in which GFP was fused to 1 kb of sequence upstream of the *sutA* open reading frame (ORF) was used to show that GFP signal strongly increased in stationary phase after growth in LB, even though mRNA levels for the reporter did not substantially change [9]. This observation suggested that post-transcriptional mechanisms might dominate control of SutA levels. However, using a combination of methods including northern blots, western blots with an antibody raised against SutA, reporter constructs, and regulator mutants, we show here that *sutA* mRNA is produced from multiple promoters regulated by an array of sigma and transcription factors, including the stress sigma factor RpoS and the flagellar sigma factor FliA. We explore dynamics of SutA expression and its direct and indirect impacts on the FliA and RpoS regulons. We find evidence that SutA is a short-lived protein whose expression increases during the transition to stationary phase, directly enhancing expression of an RpoS regulon member, but not a FliA regulon member. It contributes to both motility and RpoS-dependent extracellular proteolysis under these conditions, demonstrating that in *P. aeruginosa*, SutA is part of a starvation-responsive regulatory network that actively supports investment in resource-intensive survival-promoting activities.

## Results

### SutA levels are dynamically modulated depending on growth phase and media composition, with multiple mRNA and protein species detectable

To permit direct quantification of SutA protein levels, a polyclonal antibody against the central and C-terminal domains of SutA was raised (Fig. S8A) and western blots were performed on lysate collected across 30-50 hours of growth (starting at an optical density, OD_600_, of 0.01) in LB or phosphate-buffered minimal media with pyruvate/ammonium chloride or arginine as carbon and nitrogen sources. The growth curves differed significantly among the different media conditions and allowed identification of exponential, transition, and stationary phases (Fig. 1A-C). LB supports the fastest growth, as it is replete with vitamins, co-factors, and amino acids and small peptides that provide nitrogen and a preferred organic acid carbon source for *P. aeruginosa* [18]. Although pyruvate is a key node in central carbon metabolism, growth in pyruvate minimal media is very slow and cultures continue growing for longer than in other media, turning a vivid blue colour due to ongoing pyocyanin production (Fig. S1A). This condition was previously used to investigate SutA activity [9]. Arginine can serve as a sole carbon and nitrogen source with multiple possible catabolic pathways in *P. aeruginosa,* although ammonium chloride is a preferred nitrogen source relative to any organic source [19]. Arginine minimal media lacks the vitamins and co-factors present in LB, but supported similar growth stages, albeit with a longer lag phase and lower optical densities overall. The cells started to form clumps from T24 onwards (Fig. S1A).

**Fig 1.**
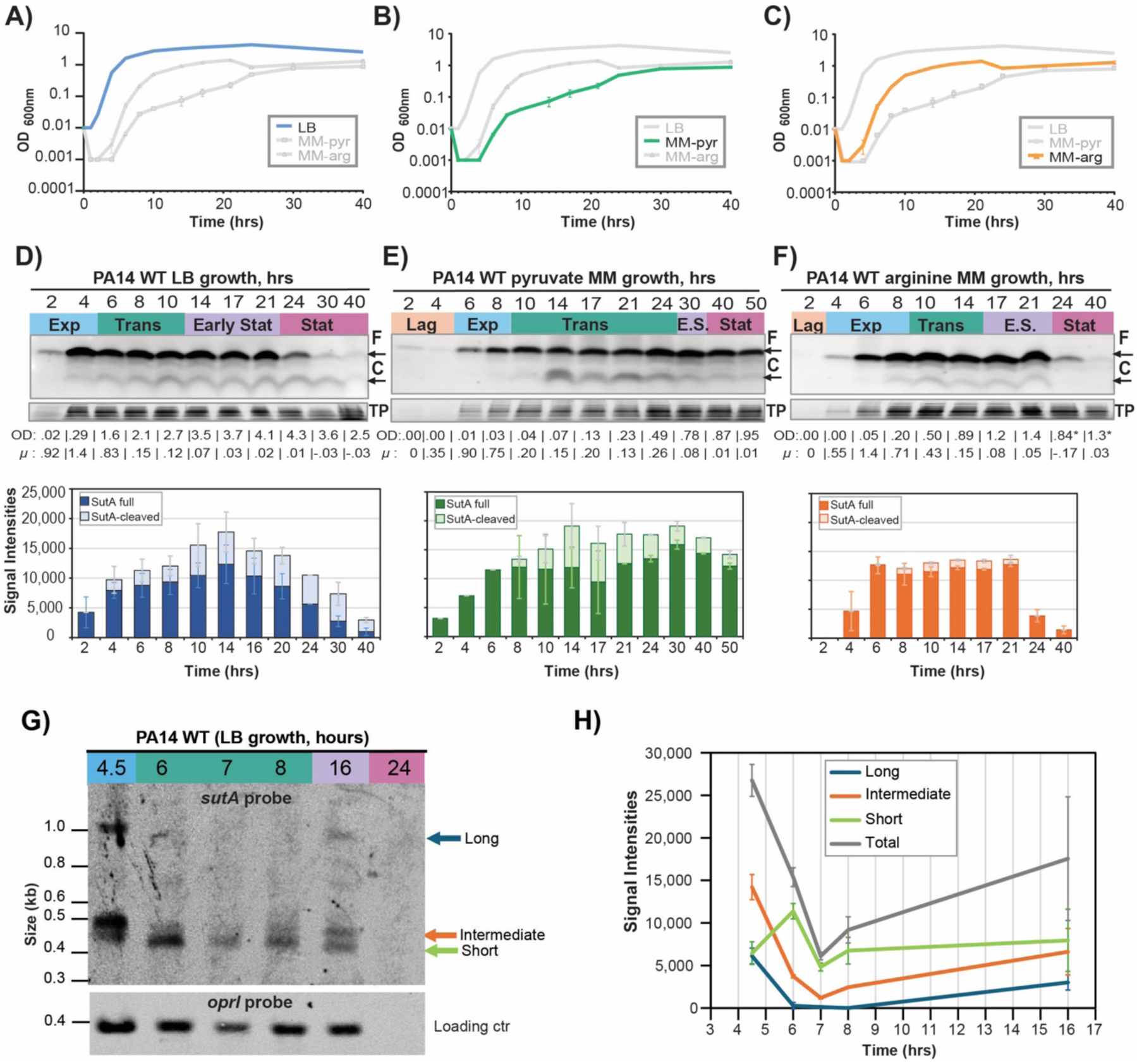
Levels of *sutA* protein and mRNA in different growth media. **A-C)** Growth curves for WT PA14 strain in 3 media. Cultures were started at OD= 0.01 in flasks and optical density of aliquots was measured by spectrophotometer at indicated times. Lines connect averages and error bars represent standard deviation from 2 independent measurements on different days and are sometimes too small to be visible. **D-F)** Western blot analysis of protein samples from different growth stages (Exp = exponential, μ>0.5); Trans = Transition, 0.1< μ<0.5; E.S.= early stationary, 0.012< μ <0.1; Stat = stationary, μ<0.012) using a polyclonal SutA antibody. Representative blots are shown at the top of each panel, alongside the OD value and the growth rate at each time point (calculated from the previous to the current time point). Quantification data for both a full-length SutA protein (11kDa; “F”) or cleaved SutA protein (∼6 kDa, “C”) are shown in the stacked bar plots below. Averages and standard deviation are shown, which are derived from at least two independent cultures and blots. All measurements were normalised by the total protein (“TP”) amounts in the gels prior to transfer. Note that the optical densities and growth rates in MM-arg cultures past T24 are unreliable due to clumping of the cells. **G)** Northern blot analysis of total RNA extracted from PA14 WT grown in LB broth at 37°C with aeration using a *sutA*-specific probe or an *oprI* probe as a loading control. A ∼1kb transcript corresponding to a possible co-transcript, as well as two smaller (∼0.4-0.5-kb) transcripts were detected. **H)** Quantification of signal intensities of all three transcripts normalised by the *oprI* loading control. Lines connect the averages and error bars show standard deviations from two independent RNA preparations and northern blots. No signal was detected in the 24 hour sample.

Western blotting with the anti-SutA antibody revealed the presence of both a full-length protein of ∼11 kDa, as well as a faint, slightly smaller protein, and a much smaller (∼6 kDa) protein, suggesting possible cleavage of SutA (Fig. 1D-F). Both the full-length and truncated versions of SutA were also observed when samples were taken four hours after transfer to minimal media without any carbon source, whereas none of these bands were detected in a *ΔsutA* mutant (Fig. S1B). The truncated versions were most abundant during the transition from rapid exponential growth phase to non-growing stationary phase all media tested. In LB, total SutA protein amounts peaked in the transition and early stationary phase and decreased later in stationary phase (Fig. 1D), whereas high levels of SutA protein were maintained for longer in pyruvate minimal medium (Fig. 1E). Total SutA levels were lowest in arginine minimal medium and rapidly reduced as soon as cultures reached stationary phase and cells started to clump (Fig. 1F), with no more cleavage product being visible. Amounts of full-length and cleaved SutA across these time courses and media conditions were quantified using ImageJ [20] (Fig. 1D-F).

We also performed western blots using an antibody against GFP in a previously generated *sutA* promoter-GFP fusion strain[9] (Fig. S1C). As previously reported, we saw that GFP accumulation continued to increase dramatically into stationary phase, a contrast to the native SutA levels that suggests differences in stability between GFP and SutA. Finally, we performed western blots against both SutA and GFP in parallel, in a strain carrying a C-terminal fusion of GFP to the full-length SutA protein, under control of its own promoter in the *attB* site of the chromosome (in addition to the native copy of *sutA*) (Fig. S1C, D). We found that this fusion protein was also cleaved, yielding a pattern of bands that was consistent with the cleavage pattern of WT SutA. Based on the observed pattern of bands, we propose that the ∼6-kDa cleavage product is the result of removing the unstructured, acidic N-terminal tail of SutA (approximately the first 40 amino acids) while the fainter, larger cleavage product is due to the removal of the C-terminal tail (approximately the last 20 amino acids) (Fig. S1E). While the mechanisms and functions of these cleavage events are outside the scope of this study, we have quantified the abundances of both the full-length SutA and 6-kDa cleavage product throughout this study, as both are evidence that SutA has been expressed.

We next performed northern blots to track *sutA* mRNA levels over the LB growth curve. Total RNA was separated on an agarose gel, transferred onto a membrane, and probed using a biotinylated RNA probe complementary to the *sutA* transcript. This revealed the presence of three distinct transcripts of ∼1-kb, ∼450-bp, and ∼400-bp in size (Fig. 1G). The long transcript and the intermediate transcript occurred in similar ratios through the growth curve, showing strong expression in exponential phase, no expression in the transition phase, and expression returning in early stationary phase (Fig. 1H). The shortest transcript was expressed in all growth phases, but peaked at T6, which coincides with a pronounced slowing of growth and the onset of pyocyanin production. We were unable to isolate enough mRNA at the 24-hour time point to assess *sutA* mRNA levels by northern blot.

### Three *sutA* transcripts can be attributed to two promoter regions and a likely post-transcriptional RNA processing event

The presence of three distinct *sutA* transcripts dynamically changing across the growth curve suggested that different regulators might drive expression in response to different environmental conditions. The bands detected by northern are consistent with a short monocistronic transcript starting directly upstream of the SutA coding sequence, an intermediate transcript starting near the end of the *PA14_69780* sequence (annotated as a putative homologue to *yjbQ*, which has no clear function[21], and henceforth referred to as *yjbQ*), and a long transcript encompassing both *sutA* and *yjbQ* coding sequences (Fig. 2A). We attempted to map the start sites of the two shorter transcripts by 5’ RACE and found a cluster of possible starts for the shortest transcript from −23 to −34 upstream of the start codon for *sutA* (Fig. S2A, B). We also obtained sequences corresponding to the intermediate transcript that mapped immediately downstream of the translational stop for *yjbQ*. Finally, we confirmed the presence of the bicistronic transcript by PCR primer walking across a cDNA anti-sense to the *sutA* transcript, and we determined that this transcript started within 40 bp upstream of the annotated translational start for *yjbQ* (Fig. S2E). There is no predicted transcription termination signal between *yjbQ* and *sutA*, and RT-PCR suggests that levels of the *yjbQ* coding sequence and the bicistronic transcript are similar while levels of the *sutA* coding sequence are higher, consistent with the existence of a bicistronic co-transcript and shorter monocistronic transcript as detected in the northern blot (Fig. S2C, D).

**Fig 2.**
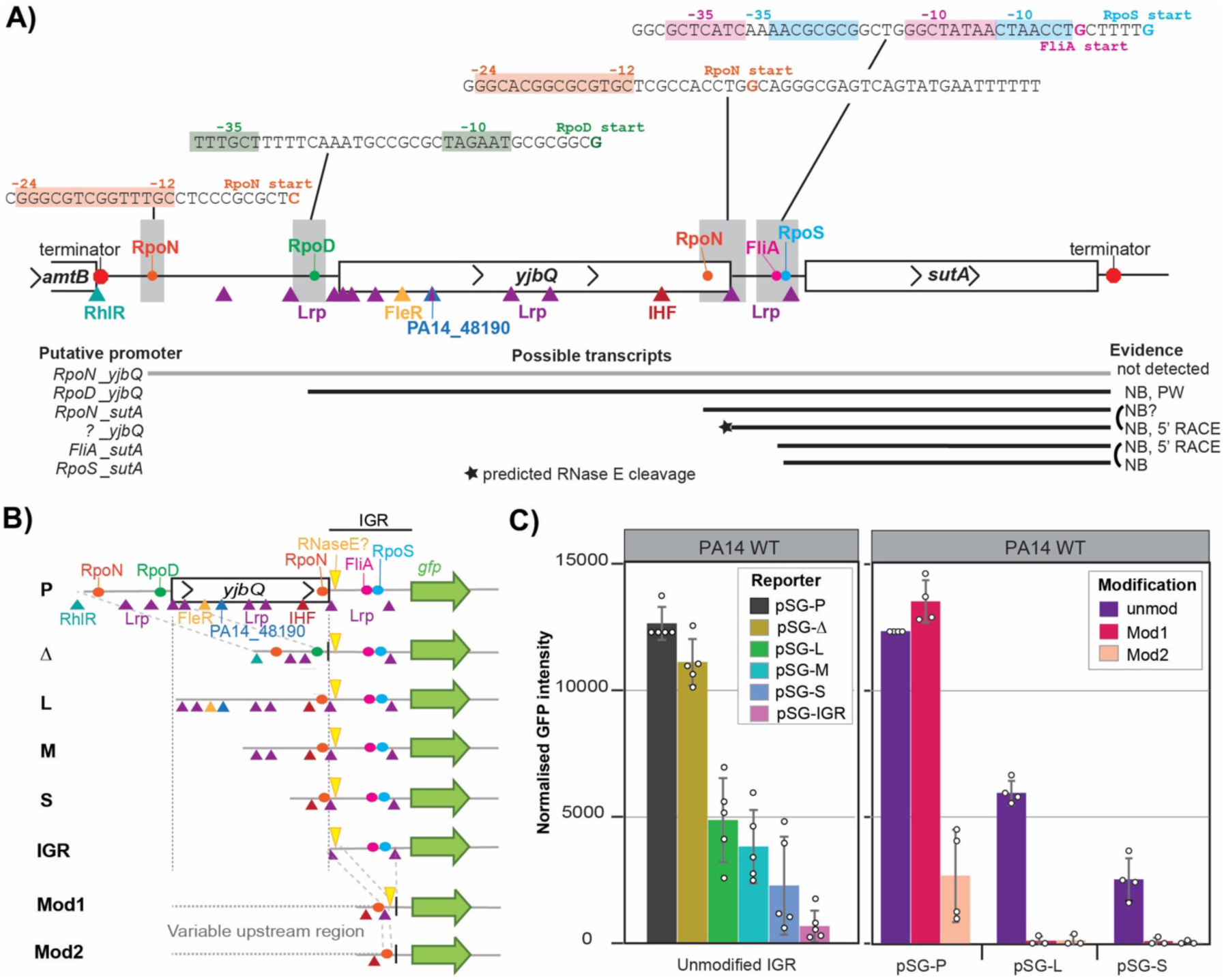
Analysis of transcription factor binding and promoter activity. **A)** Schematic of TF factor binding within the chromosomal locus surrounding the *sutA* (= *PA14_69770*) coding sequence (on negative strand in annotated genome). *In silico* predicted promoters are located upstream of *yjbQ* (=*PA14_69780*) and within the intergenic region (IGR) upstream of *sutA*. Rho-independent terminator sequences are indicated in red. **B)** Schematic of the transcriptional *sutA-gfp* reporters used in (C). The parental version (P) and the *yjbQ*-delete version Δ encode the *yjbQ* promoter(s), whereas reporters M to IGR lack them. IGR modifications were introduced into the P, L, and S variants by either deleting sequences from immediately downstream of the proposed RNaseE cleavage site up to the RBS (Mod1) or all bases immediately downstream of the translational stop codon of *yjbQ* up to the RBS, thus also deleting the proposed RNaseE cleavage site (Mod2). See Fig. S3E for sequences. The possible RpoN-dependent start within the *yjbQ* ORF is maintained in these modified reporters. **C)** Quantification of western blot samples from modified *sutA-gfp* reporters grown overnight in LB medium. Normalised, average GFP signal intensities form a monoclonal anti-GFP antibody and standard deviation from at least three biological replicates are shown. P-values for all pairwise comparisons are shown in Fig. S3D, G.

We next identified candidate factors that could drive and regulate the detected transcript starts by mining existing genome-wide transcriptomic and bioinformatic resources (Fig. 2A and Fig. S2F). Schulz *et al*. surveyed sigma factor regulons in *P. aeruginosa* using a combination of chromatin immunoprecipitation (ChIP-Seq and ChIP-chip), RNA-Seq of sigma factor deletion and overexpression strains, and searches for known consensus motifs [22]. They identified a transcription start site (TSS) for *sutA* 20 bp upstream of the start codon and called *sutA* as a member of the FliA regulon but also detected some ChIP signal for RpoS upstream of the *sutA* coding sequence. Upon inspection of the sequence, we find a strong match for the *P. aeruginosa* RpoS consensus with a TSS matching the predicted −20 site [22, 23], and a possible match for the FliA consensus with a TSS at −26 [22, 24]. This −26 TSS is supported by our 5’ RACE data, but we did not recover any transcripts matching the −20 site, possibly because the RNA for that experiment was collected during exponential phase when RpoS is not active, or due to non-templated addition of nucleotides by the reverse transcriptase [25]. The online tool SAPPHIRE, which identifies RpoD (sigma70) consensus sequences in *Pseudomonas* genomes [26], identified a high-scoring match which would drive transcription starting 32 bases upstream of the *yjbQ* start codon, consistent with the transcript boundaries suggested by primer walking experiments and the size of the long transcript in the northern blot. *yjbQ* was called as a member of the RpoN (σ^54^) regulon [22], and we could identify matches to the RpoN consensus upstream of *yjbQ* and within its coding sequence, but neither of the predicted TSS for these putative RpoN sites are supported by our northern blots and 5’ RACE data.

One possibility is that transcripts initiated by E σ^54^ are not readily detected because these transcripts are further processed. For example, no predicted sigma factor consensus sites are consistent with the intermediate transcript start, and RNaseE has been shown to be able to cleave transcripts internally at a consensus sequence of RN/WUU (with R = G/A, W = A/U, and N = any nucleotide) [27], stimulated by a nearby RNA duplex [28], which matches the context at the 5’ end of this transcript. Additionally, an Hfq cross-linking immunoprecipitation (CLIP-Seq) study identified Hfq binding sites within the ORFs of both *yjbQ* and *sutA* [29], and Hfq has been shown to direct RNaseE processing [27]. Further work will be needed to fully investigate these possible interactions.

We used the online tool PRODORIC [30] to identify putative transcription factor binding sites and found multiple matches to a consensus for the global amino acid responsive regulator Lrp, although the consensus sequence has relatively low information content so frequent matches are expected. A good match to a longer Lrp consensus defined in *E. coli* [31] overlaps with the possible RpoS −10 site upstream of *sutA*. We found matches to the consensus for the quorum sensing regulator RhlR [32] and the DNA-bending protein IHF [33] as well. We also used data from the SELEX-based transcription factor atlas for *P. aeruginosa* published by Wang *et al.* [34], which identified consensus sequences for the RpoN-dependent flagellar regulator FleR and an uncharacterised transcription factor associated with quorum sensing networks, PA14_48190 (PA1241 in the PAO1 strain).

We next carried out experiments to test the impacts of these predicted regulatory sequences. We generated a series of chromosomally encoded promoter-GFP reporter strains to examine the influence of *cis*-regulatory sequences on expression (Fig. 2B). We measured GFP in cells from overnight cultures with the aim of capturing the cumulative effects on expression throughout growth and the transition to stationary phase. First, we observed that constructs containing the upstream *yjbQ* promoter region generated substantially higher GFP signal than those lacking it (Fig. 2C and Fig. S3A-D), suggesting that the predicted RpoD-dependent promoter contributes significantly to cumulative signal over the growth curve, especially in LB medium. In minimal media, reporter levels are lower overall and less dependent on the upstream promoter. (Fig. S3C, D). We also found that signal strength dropped as increasing amounts of the *yjbQ* coding sequence were eliminated, in the absence of the upstream promoter, suggesting that these sequences contribute to supporting transcription from the proximal promoter. Finally, we investigated the impacts of removing the putative RNaseE cleavage site and the proximal promoter, in combination with some of the upstream truncations (Fig. 2C and Fig. S3E for sequences). We found that in the presence of the upstream promoter, the proximal promoter did not contribute to cumulative GFP signal, but loss of the putative RNaseE cleavage site in the Mod2 construct dramatically decreased signal (Fig. 2C and Fig. S3F). Once the upstream promoter was removed, the proximal promoter was required for any signal at all to be detected.

We also investigated whether these putative regulatory sequences were conserved across members of the Pseudomonadaceae family (Fig. S4). We found that the genomic context was conserved, with the putative transcriptional units of *sutA* found downstream of the *glnK*-*amtB* operon in all species (although in one case the *amtB* gene appears to be lost). *yjbQ* was present directly upstream of *sutA* in most but not all species (Fig. S4A). Interestingly, closer inspection of 7 representative species revealed that the upstream RpoD consensus, the putative RNaseE cleavage site, the proximal overlapping FliA and RpoS consensus, and multiple putative Lrp binding sites were conserved across all species, even if the *yjbQ* ORF had been lost (Fig. S4B). The putative RpoN consensus sites were not conserved, at least relative to the locations of the other regulatory sequences. The *glnK-amtB* operon has been shown to be regulated by RpoN in *P. aeruginosa* PAO1 and in *E. coli*, so it is possible that the wider genomic context is important for a regulatory contribution of RpoN, and our reporter constructs did not maintain this.

Together, these results suggest that during growth to stationary phase in LB, *sutA* is transcribed from an upstream promoter driven by RpoD that produces a bicistronic transcript, and a weaker proximal promoter driven by FliA and/or RpoS. The long transcript may be processed into a shorter form that is more stable and/or more competent for translation. Although we identified at least two possible RpoN consensus sequences along with IHF and FleR consensus sequences, which together could drive transcription as part of the flagellar biosynthesis cascade upstream of FliA [35], we did not obtain any evidence that these sequences were necessary or sufficient to support *sutA* GFP reporter expression under the conditions we investigated. This could be because these regulators are not active under the conditions investigated, or because our reporter constructs disrupted a required combination of sequences from different parts of the promoter region and/or the *sutA* ORF.

### Assessment of regulator mutants supports roles for RpoS and FliA at the *sutA* proximal promoter

Following our assessment of the *cis*-regulatory sequences upstream of the *sutA* ORF on expression of a GFP reporter, we next tested whether deletion or mutation of the predicted trans-acting regulators had an impact on native SutA expression. We focused on three groups of regulators: members of the flagellar regulatory cascade (FleR, FliA, and FlgM), starvation-associated regulators (RpoS and Lrp), and quorum sensing associated transcription factors (RhlR and PA14_49180). RpoD, the housekeeping sigma factor, is essential, and RpoN mutants have growth defects [36], so we did not directly investigate their contributions. We grew overnight cultures and assessed whether native SutA levels were affected by any of the mutations by western blot (Fig. 3A).

**Fig. 3:**
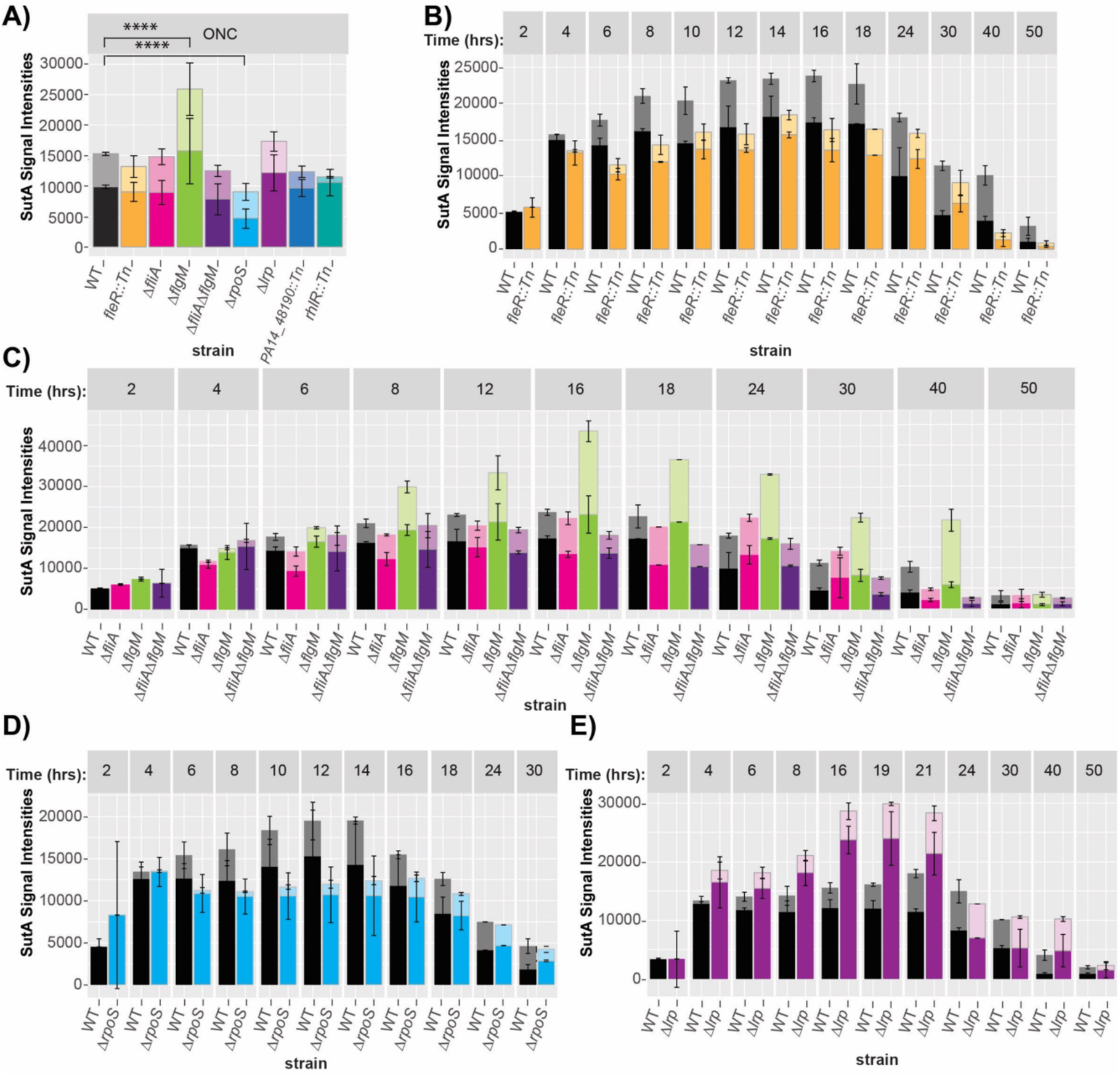
Effects of mutations on SutA expression. **A)** SutA levels in overnight cultures in LB medium of PA14 WT vs isogenic deletion mutants and transposon insertion mutants. Proteins were extracted and analysed by western blotting using a polyclonal anti-SutA antibody. Normalised SutA signal intensities (full-length = bottom and cleaved SutA = top of stacked bars) in the mutants or a WT control culture are shown as average and standard deviation from at least five independent cultures. Significant p-values for total SutA levels after a nested one-way ANOVA and a Tukey’s multiple comparisons post-test are shown. **B-E)** SutA levels in PA14 WT vs isogenic deletion mutants and transposon insertion mutants at different growth stages in LB medium. Proteins were extracted at indicated time points and analysed by western blotting using a polyclonal anti-SutA antibody. Normalised SutA signal intensities (full-length and cleaved SutA) in the mutants or a WT control culture are shown. In all graphs, the averages and standard deviations from at least two independent cultures are shown. Mutants analysed were *fleR::tn* (B), *ΔfliA* and/or *ΔflgM* (C), *ΔrpoS* (D) and *Δlrp* (E). See Fig. S5D for statistical analysis table.

Flagellar biosynthesis is tightly controlled by a hierarchical regulatory cascade. The two-component response regulator FleR, in conjunction with RpoN (σ^54^), is responsible for driving expression of class III genes including components of the flagellum membrane complex and hook. Successful construction of these complexes permits export of the anti-sigma factor FlgM, which liberates the sigma factor FliA (σ^28^ or σ^F^) to drive expression of class IV genes including flagellin (the core structural component the flagellum, encoded by *fliC*), the stator complex, components of chemosensory pathways, and potentially many other genes [22, 35, 37]. Deletion of *flgM* strongly and significantly increased SutA levels, consistent with a positive role for FliA (which is repressed by FlgM) in *sutA* expression. However, deletion of *fliA* alone or in combination with *flgM* had little effect on SutA levels, suggesting that the normal contribution of FliA under these conditions is relatively small. A transposon mutant of *fleR* showed a small (not statistically significant) reduction of SutA levels.

We observed a significant decrease in SutA levels upon *rpoS* deletion, consistent with a contribution of this sigma factor to *sutA* expression. We also observed a small (not significant) increase in SutA levels upon deletion of the gene encoding transcriptional regulator and nucleoid-associated protein Lrp. Lrp has been described as a “feast-famine response protein” and in *E. coli* can act as an activator or repressor of expression, generally activating “famine“-associated proteins such as amino acid biosynthetic pathways (necessary when exogenous amino acids are not available) and repressing “feast“-associated proteins like amino acid uptake systems (only useful when free amino acids are abundant in the environment) [38, 39]. Binding to leucine (or other free cytoplasmic amino acids) decreases its affinity for DNA, leading to derepression of genes where it acts as a repressor [40]. Our data suggest it may act in this way at the *sutA* promoter.

Disruption of the putative quorum sensing regulators *rhlR* or *PA14_48190* did not significantly impact total levels of SutA, although they did interestingly affect the fraction that was in the cleaved form. This could suggest roles in the wider regulatory network that impacts and includes SutA but does not clearly indicate a direct impact on new production of SutA so we did not follow up on these factors further.

To further interrogate the roles of the flagellar and starvation-associated regulators, we examined their impacts on SutA levels across the growth curve. Samples were collected at the indicated time points from WT and mutant cultures grown in flasks in duplicate experiments performed on different days. We again observed a modest decrease in SutA levels in the *fleR* transposon mutant throughout the transition into stationary phase (hours 6-18), and a strong decrease in late stationary phase (Fig. 3B and Fig. S5A). This could be consistent with FleR and (and thus RpoN) directly affecting *sutA* expression under some conditions, although it is also possible that loss of FleR affects *sutA* expression indirectly by affecting *fliA* activity. The *ΔflgM* mutant expresses elevated levels of SutA from hour 8 onward (Fig. 3C and Fig. S5A). The *ΔfliA* and *ΔfliA ΔflgM* mutants show slightly decreased SutA levels during the transition and early stationary phase (hours 12-18) and more strongly decreased levels in late stationary phase. Taken together, these results suggest that regulators from two levels of the flagellar regulatory hierarchy, FleR and FliA, both can contribute to expression of *sutA* during the transition and early stationary phase and even more strongly in late stationary phase.

RpoS appears to have its strongest impact on SutA levels from hours 8-14, through the transition phase (Fig. 3D and Fig. S5B). The decrease in the *ΔrpoS* mutant is consistent with a direct role as a sigma factor driving *sutA* expression, and the timing is consistent with expected activity of RpoS as growth slows during the transition to stationary phase. The *Δlrp* strain showed higher SutA levels at most time points, but the most dramatic impacts were from hour 16-21 (Fig. 3E and Fig. S5C). This is consistent with the proposed role as a transcriptional repressor of *sutA* whose DNA binding is reduced by sufficient free cytoplasmic amino acids; as amino acids become more depleted later in the transition to stationary phase, the repressive effects of Lrp could become stronger. Statistical analysis of total SutA levels throughout the growth curve by two-way RM ANOVA revealed that only the *fleR::Tn* mutant had a statistically significant effect when all time-points were considered (Fig. S5D, top panel). Shortening the time window to exponential up to early stationary phase resulted in significant effects for the *fleR::Tn, ΔfliA*, and *Δlrp* mutants, whereas only including the transition and early stationary phase gave significant effects for the *ΔfliA ΔflgM* double mutant and the *ΔrpoS* mutant, with the *ΔflgM* mutant showing significant effects throughout the stationary phase (Fig. S5D, bottom panel).

It is noteworthy that none of the mutants had a pronounced growth defect, with some of the flagella mutants even reaching higher optical densities in stationary phase (Fig. S6A). The optical densities of the mutants affecting SutA levels only began to deviate from WT at approximately the same time as the effects on SutA levels became apparent (Fig, S6B). Although some of the measured impacts on SutA levels were small, we observed that the impacts on pyocyanin production tracked as expected with the measured SutA levels (Fig. S6C). We previously observed that a *ΔsutA* strain overproduced pyocyanin, while a *sutA* overexpression strain underproduced it [9]; overnight cultures of the *ΔrpoS*, *ΔfliA*, and *ΔfliA ΔflgM* strains appeared bluer than WT, while *Δlrp* and *ΔflgM* were less blue.

Finally, we expressed two of our GFP reporter constructs in the *ΔrpoS*, *ΔfliA*, *ΔflgM*, and *Δlrp* mutants grown under the three nutrient conditions described in Figure 1 to test which *sutA* promoter might be most affected by each regulator under each condition. The reporter designated “P” included both the *yjbQ* and proximal *sutA* promoters, while the one designated “L” included only the proximal *sutA* promoter (Fig. 4A). The difference in expression between the P and L constructs was assigned to the activity of the *yjbQ* promoter. We could detect signal from both reporter constructs in all three conditions in the WT, suggesting that both promoters contributed to expression (Fig. 4B). Strikingly, expression from the proximal promoter seemed completely dependent on RpoS in LB medium, as no expression was detected from the L reporter in the *ΔrpoS* strain, while the upstream promoter was unaffected (Fig. 4C). For the flagellar mutants, the proximal promoter recapitulated the behaviour observed when measuring native SutA levels, with the *ΔfliA* mutant showing lower levels and the *ΔflgM* mutant showing higher levels (Fig. 4D). The *Δlrp* mutant did not differ significantly from the WT in reporter expression during growth in LB (Fig. 4E). Significant differences were observed only for the P construct during growth in pyruvate and arginine minimal media, but the proximal promoter appeared to contribute to these differences, as it showed smaller (though not significant) differences on its own. This could reflect the multiple predicted Lrp binding sites throughout the *sutA* upstream region. In pyruvate, *lrp* deletion caused higher levels of the GFP reporter, as observed for native SutA levels at the entry to stationary phase. In arginine minimal media, reporter expression is lower in the *lrp* mutant, as well as being lower overall (Fig. 4E). Lrp can both activate and repress individual genes in a condition-specific manner, and some of these effects may be indirect since Lrp affects hundreds of genes, including the regulator of arginine uptake in *E. coli* [38]. Together, these results suggest that the proximal *sutA* promoter region is the target of both RpoS and FliA driven transcription initiation, and that Lrp can affect both promoters, possibly repressing *sutA* expression when amino acid levels are low.

**Fig 4.**
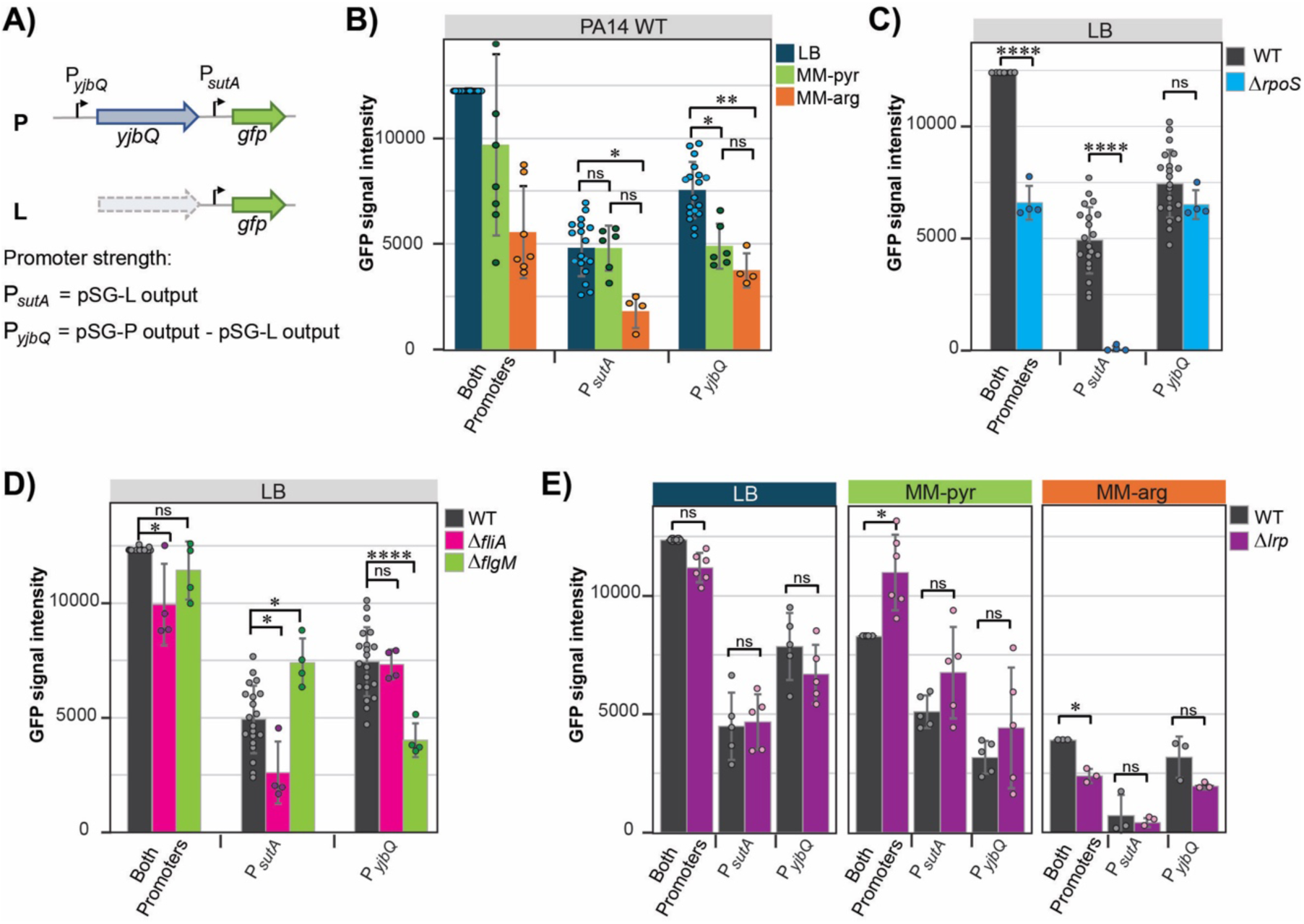
Use of *sutA-gfp* reporter constructs to assess promoter activities. **A)** Schematic of the reporters used to assess promoter activities, **B)** Promoter strength comparison using the parental vs a truncated reporter. Quantification of western blots images from overnight cultures of each reporter in either LB medium or minimal medium supplemented with either pyruvate or arginine probed with a monoclonal α-GFP antibody (representative images are shown in Fig. S3C). All intensities within a gel were normalised by Total protein (TP) visualised by stain-free technology in protein gels prior to transfer. A WT sample carrying the pSG-P reporter was included in each blot, which allowed for normalizations across multiple blots. **C)** Promoter strength comparison in PA14 WT vs *ΔrpoS* mutant reporters grown overnight in LB medium. **D)** Promoter strength comparison in PA14 WT vs *ΔfliA* or *ΔflgM* mutant reporters grown overnight in LB medium. **E)** Promoter strength comparison in PA14 WT vs *Δlrp* mutant reporters grown for 24 hrs in LB medium, or minimal medium supplemented with either pyruvate or arginine. In B-E, the P*_yjbQ_* values were calculated by subtracting the P*_sutA_* values (pSG-L) from the parental reporter (pSG-P), which contains both promoters. Averages and standard deviations from at least three blots are shown for each reporter and condition, alongside adjusted p-values after a one-way ANOVA and a Tukey’s multiple comparisons post-test.

### SutA has different impacts on RpoS and FliA regulon members

Because SutA is itself a transcriptional regulator that has previously been shown to enhance transcription initiation at rRNA promoters, especially by the Eσ^S^ holoenzyme, we wondered whether SutA might be included in the RpoS and FliA regulons so that it can help enhance transcription of other regulon members. We compared newly synthesised proteins made by the WT strain to those made by the *ΔsutA* mutant in a deep stationary phase/starvation condition using BONCAT followed by proteomics (Fig. 5A, full proteomics results in Table S1). We also incorporated RNA-Seq and ChIP-Seq data we had previously generated in pyruvate minimal media condition [9] in this analysis. For all designated RpoS and FliA regulon members (as defined by Schulz *et al.* [22]), we plotted the impact of *sutA* deletion on nascent protein and RNA levels, SutA ChIP signal, and WT RNA expression level (Fig. 5B). SutA effects on protein and RNA levels in this analysis are not expected to correlate perfectly, because the data were collected in different slow-growth conditions and because post-transcriptional regulation can play important roles in non-steady state conditions [11]. Nevertheless, the correlation was better among members of the RpoS regulon than the FliA regulon, and the average ChIP signal across all regulon members was higher for the RpoS regulon. This suggests that SutA may be more likely to directly affect transcription of the RpoS regulon than the FliA regulon.We investigated this possibility by measuring effects on transcription *in vitro* using purified Eσ^S^ and Eσ^F^ (FliA-containing) holoenzyme, prepared as previously described [15, 41] (Fig. S7C, S8B). We chose representative members of each regulon: *PA14_26020* (which we will refer to as *pepB*) [42] to represent the RpoS regulon as it was the most strongly affected by SutA and is reasonably highly expressed; and *fliC* (the most highly expressed and well-studied member) to represent the FliA regulon [24], even though our transcriptomics and proteomics data did not suggest a strong impact of SutA. We reasoned that these datasets could miss impacts if flagella were lost during collection of cells for proteomics, or if only a small subpopulation of cells were actively making flagella.

**Fig 5.**
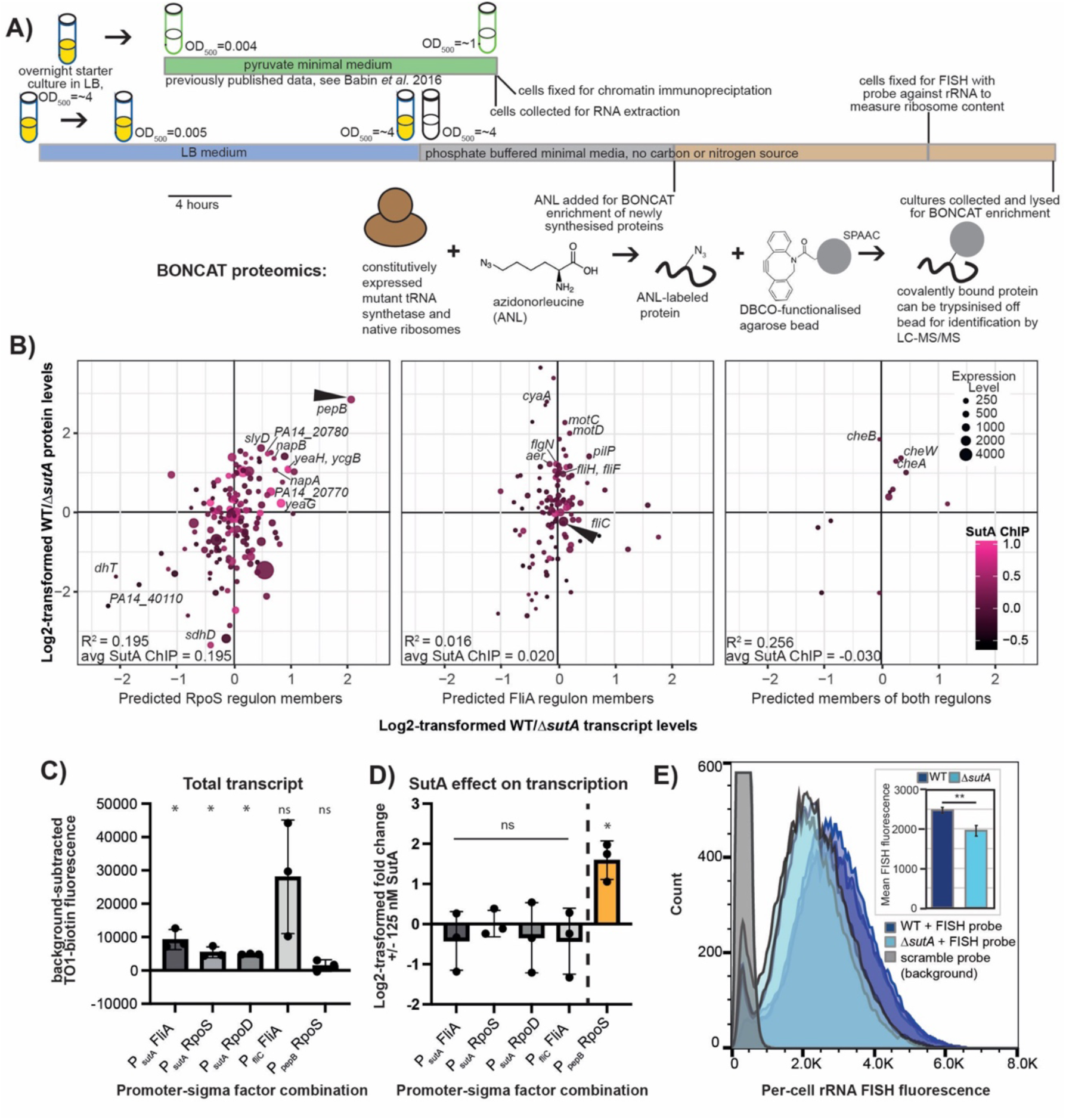
Direct and indirect impacts of SutA on RpoS and FliA regulon members. **A)** Schematic for RNA-Seq, ChIP-Seq, BONCAT proteomics, and rRNA FISH experimental time courses and BONCAT details relating to panels (B) and (E). Panel B combines previously published RNA-Seq (comparing WT and *ΔsutA* strains during late exponential phase in pyruvate minimal media) and SutA ChIP-Seq data with newly generated BONCAT proteomics data quantifying newly synthesized proteins in WT and *ΔsutA* during 24 hours of post-stationary phase incubation. SPAAC = strain-promoted azide-alkyne cycloaddition. **B)** Comparison of RNA-Seq, ChIP-Seq, and BONCAT proteomics data for members of the RpoS (left panel), FliA (middle panel) or both (right panel) regulons, as defined by Schulz *et al*. Log2-transformed fold difference in mRNA levels (WT vs *ΔsutA)* from RNA-Seq data are shown on the x-axis; log2-transformed fold difference in new protein synthesis from BONCAT-proteomics data (WT vs *ΔsutA)* are shown on the y-axis; total transcript abundance in RPKM from the WT strain are indicated by the size of the point, and SutA ChIP signal is indicated by the colour scale for each point. Genes of interest potentially affected by SutA either at the level of transcription or translation (via the potential for increased ribosome biogenesis) are annotated. **C)** *In vitro* transcription assays to measure transcription by Eσ^F^, Eσ^S^, and Eσ^70^ from *sutA, fliC* and *pepB* promoters fused to the sequence for the fluorogenic RNA aptamer mango II (see materials and methods). n=3. One-sample t-tests were used to assess whether signal was significantly different from zero. Background-subtracted Eσ^F^ activity was variable but always higher than for other holoenzymes. Negligible signal above background was detected for the *pepB* promoter. **D)** *In vitro* transcription assays were used to measure direct impacts of SutA on transcription initiation by Eσ^F^, Eσ^S^, or Eσ^70^ from the *sutA* promoter, Eσ^F^ from the *fliC* promoter, and Eσ^S^ from the *pepB* promoter. For all combinations except Eσ^S^ /*pepB* promoter, assays were carried out in triplicate as for (C), with the addition of either 125 nM SutA or an equivalent volume of storage buffer during the open complex formation step. For Eσ^S^ /*pepB* promoter (orange bar), because we could not reliably detect signal above background using the mango II reporter, a gel-based assay was used (n=3) (see materials and methods, Fig. S7A). Band intensities were quantified using the ImageJ gel lane tool. One-sample t-tests were used to assess whether the log2-transformed ratios were significantly different than zero. **E)** Fluorescence *in-situ* hybridisation with oligonucleotide probes against the ribosomal RNA, followed by flow cytometry, was used to quantify per-cell ribosome abundance in WT and *ΔsutA.* A population of likely single cells was identified by gating on DAPI signal and forward scatter, and the peak height of Cy5 fluorescence was taken as a measure of ribosome abundance for each event. The grey histogram indicates representative background, dark blue is WT rRNA signal and light blue is *ΔsutA* rRNA signal. Inset bar chart displays the mean values of background-subtracted population means for biological triplicates. P-values were calculated using an unpaired t-test with Welch’s correction.

We generated linear double stranded DNA templates containing promoters and the initial coding sequence for *pepB* and *fliC* fused to the fluorogenic mango II aptamer sequence [16, 43]. We also generated a template in which the proximal promoter and coding sequence of *sutA* was fused to mango II. We first verified that Eσ^S^ and Eσ^F^ can initiate transcription at the proximal *sutA* promoter, as our *in vivo* analyses suggested should be possible. We saw that both Eσ^S^ and Eσ^F^ drove similar or higher levels of expression than the housekeeping holoenzyme Eσ^70^(Fig. 5C). Although Eσ^S^ and Eσ^70^ recognise similar consensus motifs, Eσ^70^ should be able to drive much higher levels of expression from *bona fide* housekeeping promoters [17], so we conclude that the *sutA* proximal promoter favours Eσ^S^ over Eσ^70^.

We next asked whether SutA could directly influence transcription by Eσ^S^ or Eσ^F^ on promoters from their regulons, including the *sutA* promoter. We found that inclusion of SutA at 125 nM, which we previously showed could upregulate rRNA transcription [15] had no significant impact on transcription initiation of *sutA* by either holoenzyme or on initiation of *fliC* by Eσ^F^ (Fig. 5D). For the *pepB* promoter, we did not obtain sufficient signal from the mango II aptamer, so we used a traditional single turnover initiation assay in which transcripts are detected by incorporation of radiolabelled CTP, as previously described [15]. For this promoter, we observed a 2–4-fold increase in run-off transcript when SutA was added to the reaction at 125 nM (Fig. 5D and Fig. S7A). These data suggest that SutA can enhance transcription initiation at the promoter of an RpoS regulon member (*pepB*) but it does not generally increase initiation activity by Eσ^S^ or Eσ^F^ on all promoters they recognise.

An alternative hypothesis is that SutA may be included in the FliA regulon because its previously reported stimulation of ribosome-associated genes contributes to flagellar biosynthesis by ensuring the maintenance of sufficient protein synthesis capacity to build a flagellum, even under challenging conditions. To investigate this possibility, we used fluorescence *in situ* hybridisation (FISH) with an rRNA-directed probe to estimate per-cell abundance of ribosomes in WT and *ΔsutA* cells during the late stationary phase/starvation condition in which we performed the BONCAT proteomics experiment (Fig. 5A). We found that WT cells showed slightly but significantly higher levels of FISH fluorescence than *ΔsutA* cells (Fig. 5E and Fig. S7B). Therefore, even if SutA does not directly support transcription initiation at an abundantly expressed flagellar gene like *fliC*, it could support flagellar biosynthesis by helping to ensure that sufficient cellular resources are invested in ribosomes, under starvation conditions.

### Per-cell reporter levels indicate uniform but bursty expression of *sutA, pepB,* and *fliC* during the transition to stationary phase

We wanted to directly examine per cell expression dynamics for *sutA*, *pepB*, and *fliC* during the transition into stationary phase. Previous work measuring per-cell transcript levels for 105 genes in *P. aeruginosa* suggested that on average, both *fliC* and *rpoS* are among the most abundantly produced mRNAs during stationary phase, but only a small minority of cells contain detectable transcript at a given sampling time [44, 45]. Another study in *Salmonella enterica* detected a bimodal distribution of ribosome abundances among cells during the entry to stationary phase [46]. We wondered whether SutA might contribute to population heterogeneity during the transition to stationary phase, integrating multiple regulatory signals to support new protein synthesis in only a subpopulation of cells. Measuring single-cell expression dynamics during the transition to stationary phase is complicated by the generally low activity levels at the entry to growth arrest, and accumulation of reporter signal from highly stable reporter proteins like GFP that may have been expressed much earlier in growth. GFP reporter signal can dynamically increase and decrease in growing populations of cells due to its dilution by growth after downregulation of transcription initiation, but as growth slows and stops, any expressed GFP will accumulate, obscuring ongoing transcriptional dynamics.

We generated reporter constructs consisting of the promoter and initial coding sequences of *sutA* (equivalent to the pSG-P promoter region in Fig. 2), *pepB*, and *fliC* fused to the HaloTag sequence [47], and integrated them into the chromosomes of both the WT and *ΔsutA* strains at the *attB* site [48]. HaloTag is a modified dehalogenase enzyme that can covalently attach a chloroalkane fluorescent ligand to itself, retaining it at the site of expression of the HaloTag and increasing its fluorescence [47]. Importantly, the HaloTag is also capable of modifying itself with a chloroalkane ligand that is not fluorescent, thus blocking its ability to subsequently be modified by a fluorescent ligand [49] (Fig. 6A). We used the “dark” ligand 6-chlorohexanol (6-CHO) to quench pre-existing HaloTag reporter at the 14 hr timepoint during the transition to stationary phase, and then measured subsequent new synthesis of the reporter by washing away 6-CHO and introducing the brightly fluorescent ligand JF646 for either 15 minutes (0 hours time point), 4 hours, or 8 hours (Fig. 6A). Live cells were washed to remove excess ligand, and fluorescence was visualized by microscopy.

**Fig. 6.**
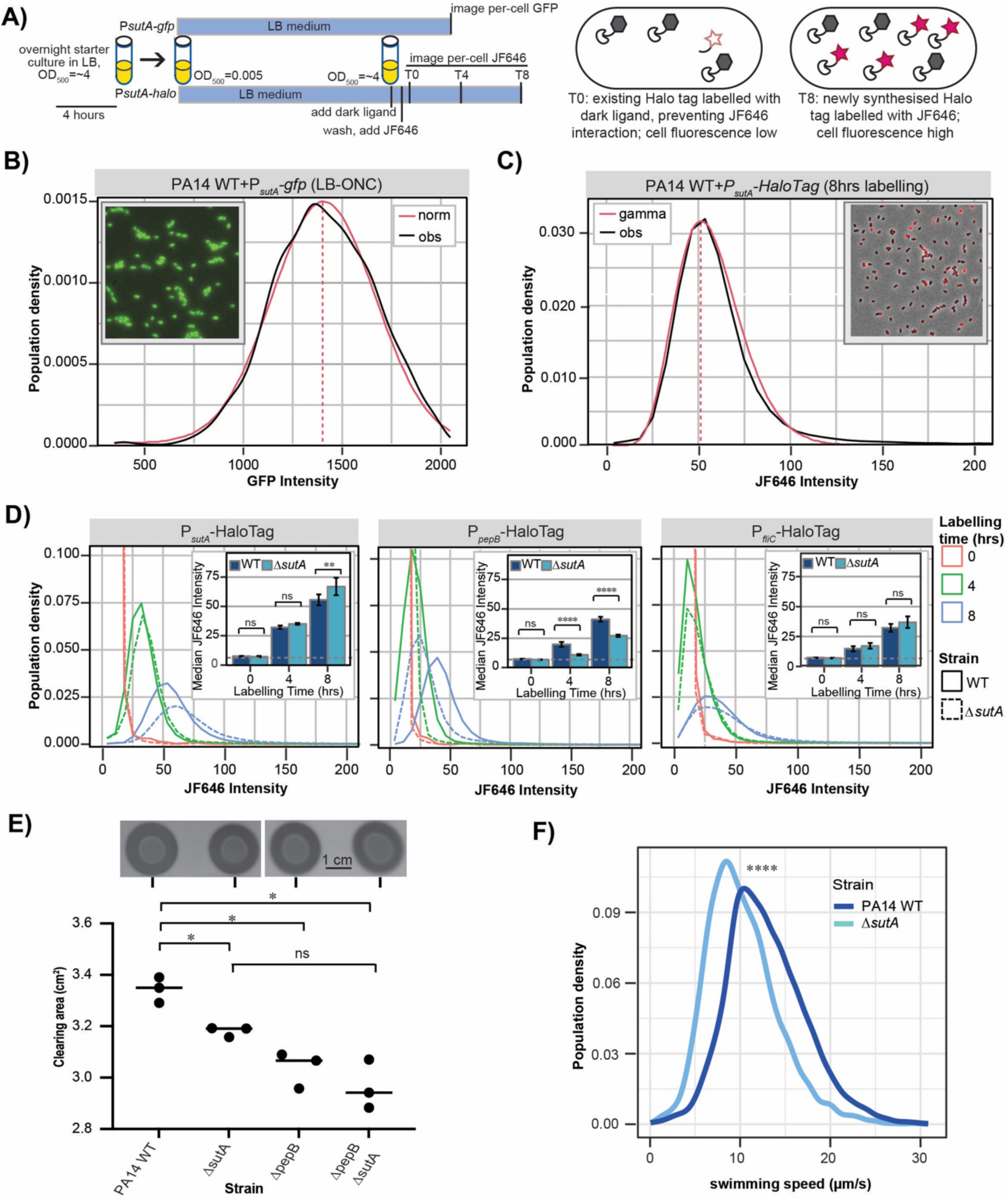
Dynamics and impacts of *sutA* expression in stationary phase. **A)** Schematic explaining HaloTag and GFP reporter experimental time courses. **B)** Distribution of per-cell fluorescence for WT cells carrying the *sutA*-GFP reporter (pSG-P version). The empirical distribution (black) approximates a normal distribution (red). **C)** Distribution of per-cell fluorescence for WT cells carrying the HaloTag fused to the pSG-P promoter region, which allows detection of new expression from the *sutA* promoters during hours 14-22 of the growth curve. The empirical distribution (black) was left-skewed and fit to a gamma distribution (red). **D)** Per-cell HaloTag fluorescence distributions for HaloTag reporters under the control of *sutA* (left panel), *pepB* (middle panel), or *fliC* (right panel) promoters, in WT (solid lines) and *τιsutA* (dashed lines) strain backgrounds, labelled for 0 (red), 4 (green), or 8 (blue) hours after quenching existing HaloTag protein with the 6-CHO dark ligand. Insets contain bar graphs showing means of population median values for three biological replicates, with error bars showing standard deviations. Significance of differences bewteen WT and *τιsutA* strains for each timepoint and promoter construct was assessed by performing a one-way ANOVA with Tukey’s post-hoc test for multiple comparisons. **E)** Secreted protease activity was assessed by spotting equivalent OD units of cells onto milk plates and allowing growth for 18 hours. The area of the halo surrounding each spot was measured using ImageJ, averaging 3-4 technical replicates for each of 3 biological replicates. Data points represent biological replicates. Significance of differences among strains was assessed using a one-way ANOVA with Dunnett’s T3 post-hoc test for pairwise comparisons. **F)** Measurements of swimming speeds of WT and *τιsutA* cells after 15 hours of growth in LB. Timelapse phase contrast microscopy images were captured at a rate of 3 frames per second and the TrackMate plug-in of ImageJ was used track individual cells and calculate their average speeds. Swimming speed data was collected from six biological replicates over two separate days. Population density plots show information from n=4742 individual cell tracks for WT and n=4328 cell tracks for *ΔsutA.* A p-value was calculated using a two-way t-test with Welch’s correction.

The *sutA*-GFP reporter (pSG-P) displayed a population distribution of per-cell GFP fluorescence that fit closely to a normal distribution (Fig. 6B). However, as shown in Fig. 2, this signal is dominated by expression from the upstream RpoD promoter, which is most active during exponential phase. In contrast, we observed that the distribution of per-cell fluorescence in the WT for the P*_sutA_*-HaloTag reporter was positively skewed and had a higher kurtosis than a normal distribution (Fig. 6C). In growing *E. coli*, stochastic bursts of reporter expression (which are seen for most promoters) have been reported to result in skewed distributions of per-cell reporter signal when the number of bursts per cell division time was small. These distributions could be fit to gamma distributions, with the number of expression bursts per division proportional to the shape parameter alpha [50]; the gamma distribution approximates a normal distribution if the shape parameter is large. The skewness and kurtosis of the per-cell HaloTag signal distributions could suggest that limiting our detection to new expression during the 14-22 hour time frame in the growth curve allowed us to detect a relatively small number of bursts of expression, while the approximately normal GFP reporter distribution is consistent with accumulation of a larger number of bursts of expression.

A comparison of per-cell HaloTag reporter fluorescence distributions at 0, 4, and 8 hours of labelling, in the WT and *ΔsutA* backgrounds, further supports that expression of *sutA* occurs discontinuously in individual cells during the entry into stationary phase (Fig. 6D). In the 4-hr labelling, some cells still showed no signal above the 0-hr background distribution, but by 8 hours most cells had some fluorescence and the distribution was broader. Similar patterns were seen for the *pepB* and *fliC* HaloTag reporters, with all the 4-hr and 8-hr distributions showing skewness and kurtosis associated with a gamma distribution with relatively low shape parameter (Fig. S9 and Fig. S10). Only the *pepB* reporter showed a significant positive effect of SutA, and the signal from the *sutA* HaloTag reporter was significantly repressed by the presence of SutA at the 8 hr timepoint, while the *fliC* reporter was not significantly affected by SutA.

We next tested whether SutA contributed to phenotypes dependent on successful expression of RpoS and FliA regulon members. *pepB* encodes a secreted aminopeptidase (also known as PaAP) controlled by RpoS, quorum sensing, and SutA (as demonstrated above) [42, 51]. We visualised activity of secreted proteases and peptidases by spotting equivalent amounts of wt, *ΔsutA*, *ΔpepB*, and *ΔsutA ΔpepB* strains on solid media containing 1.5% agar and 1.5% non-fat dry milk (w/v). After 18 hours, hydrolysis of the milk casein by secreted proteases and peptidases resulted in a zone of clearing around the bacterial spots. We observed that the *ΔsutA*, *ΔpepB*, and *ΔsutA ΔpepB* strains showed small but significant decreases in the areas of their zones of clearing compared to WT (Fig. 6E). Because multiple proteases controlled by different regulatory processes can contribute to casein hydrolysis, the defect is not expected to be complete.

To measure the outcomes of flagellar biosynthesis regulation during the transition to stationary phase we measured swimming speeds of individual bacteria directly by timelapse microscopy, after approximately 15 hours of growth in LB. Standard swimming assays, in which the area of expansion of an inoculum in soft agar is measured, could not easily be adapted to investigating stationary phase motility, because these assays require bacterial growth. We plotted the distributions of swimming speeds for WT and *ΔsutA* cells from six independent cultures grown on two different days. We observed a modest shift to slower swimming speeds in the *ΔsutA* mutant compared to the WT (Fig. 6F). Thus, SutA can contribute to optimal flagellar function during the transition to stationary phase, even if it does not directly or indirectly enhance expression of *fliC*, the most abundant flagellar component, under this condition.

## Discussion

We previously showed that SutA is a regulator expressed during slow growth and growth arrest in *P. aeruginosa* that can stimulate expression of biosynthetic machinery including ribosomal components. Here we have identified regulators that impact SutA expression to gain insight into signals associated with new protein synthesis in a context of nutrient limitation (Fig. 7A). Our results suggest that *sutA* expression as part of a bicistronic mRNA is driven by the housekeeping sigma factor RpoD during exponential growth, but that the flagellar regulators FliA, FlgM, and FleR, and starvation-associated regulators RpoS and Lrp control transcription of a monocistronic mRNA during the entry to stationary phase. Our results have also highlighted that SutA is likely to be short-lived, produced by and helping to facilitate short bursts of new protein synthesis to aid survival without overtaxing available resources.

**Fig. 7.**
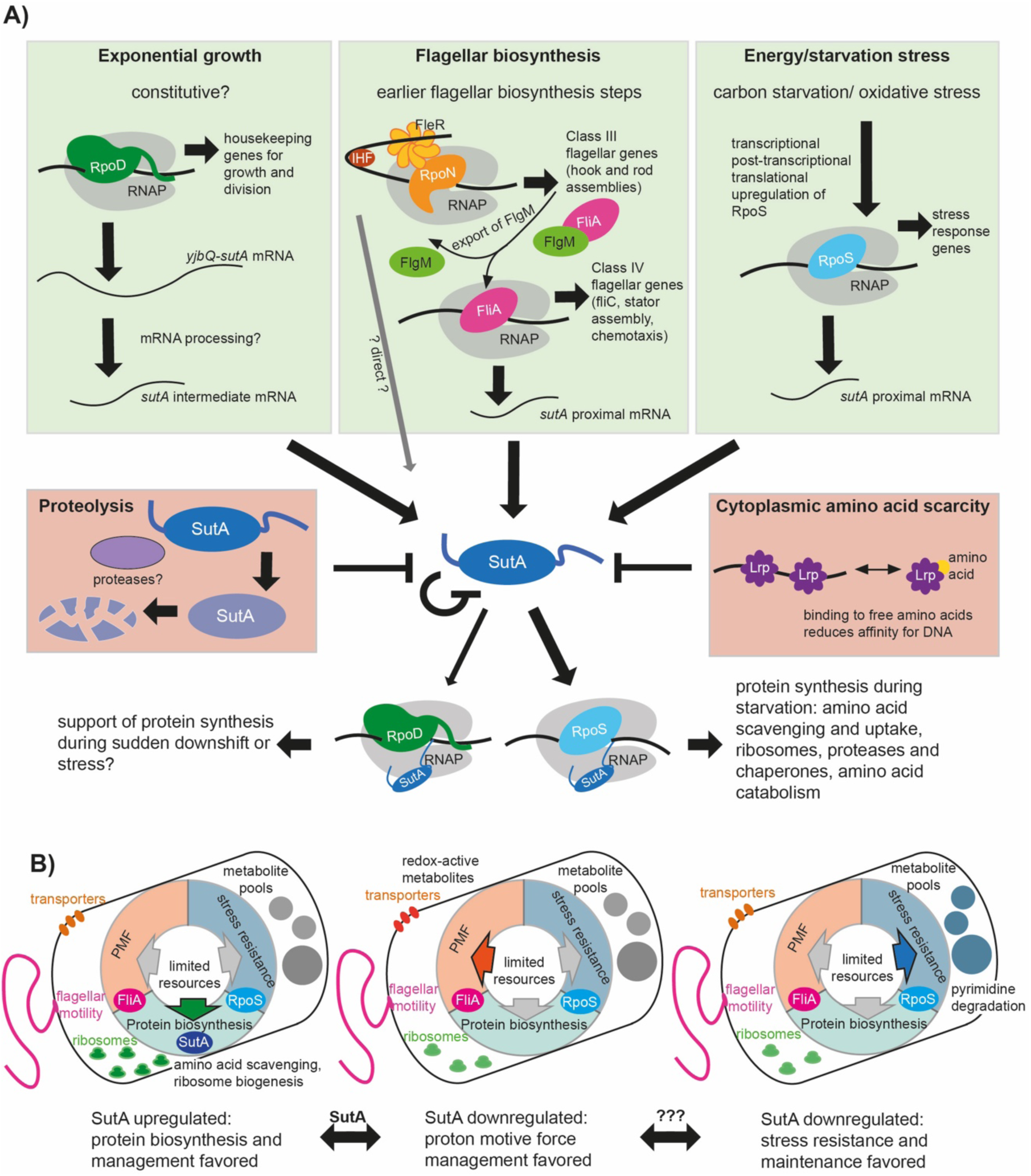
A model of SutA regulation and impacts during scavenging. **A)** Regulators identified in this work to affect SutA levels. The top row of green boxes depicts sigma factors that directly drive SutA transcription, and the signals or conditions under which they are active. The red boxes depict factors or processes that repress SutA levels. SutA appears to repress its own production, although this may be indirect. SutA has been shown to affect transcription by both Eσ70 and EσS, although the impact is larger for EσS. Constitutive expression during growth may ensure that some SutA is available in the case of a sudden nutrient downshift. **B)** Role of SutA in the wider regulatory network. SutA appears to favour expression of genes associated with protein synthesis, pushing the cell to devote resources to biosynthesis. During scavenging, this could be important for accessing, taking up, and incorporating amino acids from diverse but potentially non-preferred environmental sources. In the absence of SutA (as evidenced by mutant phenotypes), more resources are devoted to management of the PMF and stored metabolite pools (see also Table S1).

### Scavenging activities during starvation

The identification of these regulators suggests that SutA is part of a network controlling scavenging behaviours upon nutrient downshift. We define scavenging as an active search for new nutrient sources, involving motility (controlled by FliA, FlgM, and FleR), secretion of extracellular enzymes like PaAP for degradation of complex nutrient sources (controlled by RpoS), and uptake of liberated amino acids (controlled by Lrp). Many heterotrophic Proteobacteria engage in cycles of colonising organic particles or plant and animal hosts, then surviving periods of low nutrient availability while searching for a new niche after dispersal. This search is an energetically expensive gamble, and many individual cells will not be successful. However, the rewards for success can be large, and a recent modelling study has suggested that only a small fraction of starved, searching cells need to be successful for this strategy to maintain population growth [52]. Searching while starving requires continual depletion of internal stores [6], and a scavenging regulatory strategy must partition these limited resources among essential processes such as making new proteins, maintaining the proton motive force (PMF) that powers flagellar motility and transport, and preserving metabolite pools (Fig. 7B). We propose that SutA tips the balance toward new protein synthesis.

In *E. coli* during nutrient downshift, FliA first drives expensive flagellar biosynthesis to facilitate the search for new resources, and then RpoS controls later gene expression as priorities shift to preserving resources and carrying out essential stress responses [11]. Counterintuitively, mutations in both RpoS and Lrp give a growth advantage during a protracted stationary phase, where slow growth is fuelled by scavenging from dead cells [53, 54]. However, this is consistent with the notion that these regulators can restrain scavenging activity to preserve resources. Complex regulatory connections among flagellar biosynthesis, starvation responses, and protein biosynthesis suggest ongoing crosstalk may be important in naturally fluctuating conditions. For example, Lrp can indirectly stimulate expression of flagellar genes, and its impact on amino acid transporter expression depends on amino acid availability [55]. A regulator called FliZ, co-expressed with FliA, can directly repress transcription of some RpoS regulon members [56]. Recent work has also identified a regulatory connection between FliA-transcribed sRNAs and production of the ribosomal protein S10, which plays a second role as a member of a transcription anti-termination complex. This suggests that FliA-dependent modulation of the core gene expression machinery may help the cell to manage the cost of flagellar biosynthesis in nutrient-limited conditions [57].

In *P. aeruginosa*, FliA, RpoS, and Lrp are conserved but many of the modulators of their expression and activity are not [58], and the regulatory architecture achieves somewhat different outcomes. Our results suggest that SutA modulates gene expression during nutrient limitation as part of both the FliA and RpoS regulons. In a departure from the *E. coli* paradigm, the impacts of FliA and RpoS on SutA expression appear to be temporally overlapping, with *ΔfliA*, *ΔflgM*, and *ΔrpoS* all impacting population SutA levels from hours 8-16 in our LB time course. Interestingly, the flagellar regulators also appear to have an influence very late in stationary phase in our time course (Fig. 3B, C). These temporal differences in flagellar and RpoS-driven regulation compared to *E. coli* could be consistent with the observations that *P. aeruginosa* maintains flagellar motility and robust new protein synthesis deeper into starvation [5, 6]. Also different from *E. coli*, *P. aeruginosa* produce only a single polar flagellum, and cryo-electron tomography studies have suggested that it can be ejected during starvation to save resources [59]. Regulated flagellar loss could be a reason why ongoing new flagellar synthesis might be needed even in the absence of cell division. These differences suggest perhaps an even greater need in *P. aeruginosa* for coordinating resource usage among new protein synthesis, PMF management, and resource maintenance during scavenging. While not homologous to the *E. coli* system, SutA could play an analogous role in *P. aeruginosa* to the FliA-dependent S10-modulating sRNAs by tuning abundance and activity of the core gene expression machinery.

### Gene expression dynamics during growth arrest

Our investigation of *sutA* regulation during the entry into stationary phase has illuminated some general issues surrounding regulatory paradigms outside of steady state growth. First, the use of fluorescent proteins as reporters of gene expression is complicated by their stability when the proteome is not being diluted by growth. GFP has an estimated half-life of more than a day [60], so it can serve as a reporter of cumulative production but changes to expression occurring after nutrient depletion, when total activity is low, might be obscured. Second, low overall activity during growth arrest imposes a limit on the magnitudes of regulatory changes that can be achieved. Biosynthetic activity must be tightly controlled to limit the resource costs of engaging in any protein production, and fine-tuning by post-transcriptional regulatory mechanisms may play a more important role than during exponential growth, where transcription initiation controls levels of most proteins [61]. Third, population heterogeneity may play important roles in growth arrest, allowing a subset of cells to engage in an expensive activity that could benefit the wider population. This activity could be difficult to detect at the bulk population level.

Although we did make use of GFP reporters to test cis-regulatory sequence impacts on cumulative SutA production, we sought to overcome the difficulties in detecting per-cell new expression dynamics during the entry to stationary phase by using a HaloTag reporter. The JF646 fluorophore ligand we used is very bright, with a higher quantum efficiency and almost 10-fold higher molar extinction coefficient than the fluorescent protein PA-mCherry [62]. A similar system was reported to detect 7 copies per cell of the HaloTag reporter in *E. coli* [63]. This permitted us to focus only on reporter that was newly synthesised during the entry to stationary phase, after existing reporter molecules were quenched with a dark ligand. We observed a large difference in fluorescence between quenched and unquenched control samples at the 0-hr time point as expected, but this difference diminished at the 4- and 8-hr time points, which suggests that the Halo Tag reporter may also be substantially less stable than GFP (Fig. S9A).

We did not observe any evidence of a bimodal distribution, with a highly active subpopulation, in the three reporters we constructed. Instead, we could fit gamma distributions with relatively low shape parameters (alpha = 3-12) to all distributions. According to previous work fitting per-cell expression data to gamma distributions [50], these distributions could be explained by the stochastic bursts of expression proposed to characterise most genes during growth, but with relatively few bursts during the long labelling times. The different reporters produced distinct distributions, with the *fliC* reporter displaying a broad distribution with a high scale parameter potentially indicating larger bursts of expression in this model. This could be consistent with the previously reported mRNA FISH results showing that *fliC* transcripts were highly abundant in a small fraction of the stationary phase cells of *P. aeruginosa* [44, 45]. Because mRNAs are very short-lived, bursty expression and a large burst size could produce this pattern; our data suggest that most cells experience at least one such burst of *fliC* transcription at some point over 8 hours at the entry to stationary phase. We do not know the HaloTag reporter’s half-life or the amount of signal per reporter molecule, and we measured signal accumulated over several hours, so we cannot extract absolutely quantitative information about the expression burst numbers or sizes from our per-cell signal distributions as was done previously for reporter expression in *E. coli* [50]. Further characterisation of expression dynamics during growth arrest will be an interesting area for future work.

Our data suggest a unimodal distribution of *sutA* expression across the population of planktonic cells at the entry to stationary phase, but the complex promoter architecture and post-transcriptional and post-translational controls on SutA levels suggest that multiple signals can be integrated to permit sensitive responses to environmental conditions. Several observations also suggest that SutA is likely to be unstable: 1. We detect direct evidence of multiple cleavage products in western blots; 2. Our GFP reporter levels suggest a high contribution of an RpoD-driven exponential phase promoter and low contributions of an RpoS-and FliA-driven transition phase promoter to cumulative levels of GFP, but total native levels detected by western are highest during the transition into stationary phase, suggesting SutA produced in exponential phase has been turned over; 3. Previously published proteomic data comparing the newly synthesised (BONCAT) proteome to the total proteome in carbon starvation showed that SutA occupies a nearly 10-fold higher fraction of the newly synthesised proteome, which can be explained by high turnover [7]. High turnover of a stationary phase regulator could allow its levels to be temporally dynamic despite a lack of proteome dilution by growth. More work will be required to understand the possible inputs of RpoN, FleR, Hfq, RNaseE, and yet unidentified protease(s) to controlling levels of SutA. Dynamic and environmentally responsive expression of SutA, a facilitator of new protein synthesis, can support active survival strategies within constraints imposed by limited resources. These strategies likely contribute to the ecological successes of *P. aeruginosa* in a wide range of environments.

## Materials and Methods

### Bacterial Strains and growth conditions

All bacterial strains used in this study are listed in Supplemental Table S2. Bacterial overnight cultures were grown in 5 ml volumes of media in universal tubes with loosened lids. Growth curve cultures were grown in 50 ml volumes of media in 250-ml conical flasks. All cultures were grown at 37°C with shaking at 200 rpm. Liquid media were LB (5 g/L yeast extract, 10 g/L tryptone, and 10 g/L NaCl or 5 g/L for low-salt version) or phosphate-buffered minimal medium (35.9 mM K_2_HPO_4_, 14.2 mM KH_2_PO_4_, 42.8 mM NaCl, 1.0 mM MgSO_4_, 7.5 µM FeCl_2_·4H_2_O, 0.8 µM CoCl_2_·6H_2_O, 0.5 µM MnCl_2_·4H_2_O, 0.5 µM ZnCl_2_, 0.2 µM Na_2_MoO_4_·2H_2_O, 0.1 µM NiCl_2_·6H_2_O, 0.1 µM H_3_BO_3_, and 0.01 µM CuCl_2_·2H_2_O) with carbon sources added at a 20 mM concentration and 9.3 mM NH_4_Cl as nitrogen source, where applicable.

### Construction and verification of unmarked deletion or transposon mutants in UCBPP-PA14

All unmarked deletion mutants were generated by PCR amplifying 500 bp of the upstream and downstream sequences of each target gene using Q5 2x Master Mix (New England Biolabs, Cat # M0492S) and genomic DNA of UCBPP-PA14 as a template, and ligating and cloning the PCR products into linearised suicide vector pMQ30 (using enzyme EcoRI and HindIII, both New England Biolabs, Cat # R3101S and R3104S, respectively) by Gibson assembly using the NEBuilder HiFi Assembly Mix (New England Biolabs Cat # E2621S). Reactions were transformed into *E. coli* DH5α cells by electroporation and all constructs were verified by Sanger sequencing. The constructs were subsequently introduced into *P. aeruginosa* UCBPP-PA14 by triparental conjugation using strain HB101/pRK2012 as a helper strain. Successful exoconjugants were selected on VBMM medium (3 g/L trisodium citrate, 2 g/L citric acid, 10 g/L K_2_HPO_4_, 3.5 g/L NaNH_4_PO_4_, 1 mM MgSO_4_, 100 μM CaCL_2_, pH 7) containing 100 μg/mL gentamicin as described by Choi and Schweizer [48] and were then subjected to counterselection on LB plates lacking NaCl but containing 10% (w/v) sucrose. Colonies resulting from homologous recombination to remove the wild-type copy of each target gene and retain the clean deletion were identified by PCR and confirmed by whole genome sequencing (Plasmidsaurus). Transposon mutants were retrieved from the ordered non-redundant UCBPP-PA14 transposon library[64], PCR verified and confirmed by whole genome sequencing. Genome sequencing was analyzed to confirm the absence of unexpected SNPs using the breseq package[65].

### Construction of reporter strains in UCBPP-PA14

Modified versions of a *sutA-gfp* transcriptional fusion reporter plasmid pUC18T-mini-Tn7T-Gm^R^-P*_sutA_:sfgfp* (extracted from strain DKN1641, see Babin *et al* 2016 [9]) were created by amplifying smaller parts of the plasmid using relevant fw and rv primers (see Table S2) with and without modified linkers using Q5 2x Master Mix (New England Biolabs, Cat # M0492S) and recircularizing the plasmids by Gibson assembly using the NEBuilder HiFi Assembly Mix (New England Biolabs Cat # E2621S). Other reporter constructs were also made by Gibson assembly. Mango II reporters were constructed by assembling the relevant promoter and initial or complete coding sequences amplified from the PA14 genome (using Q5 2x Master Mix), the mangoII aptamer sequence amplified from plasmid pLenti-mCherry-Mango II x 24 (using KOD One 2x hot start master mix (Sigma)), and the pUC18T-mini-Tn7T-Gm^R^ backbone, using primers indicated in Table S2. HaloTag reporters were constructed by assembling the relevant promoter and initial coding sequences from the PA14 genome, the HaloTag sequence from plasmid pET51b-His-TEV-HaloTag7, and the pUC18T-mini-Tn7T-Gm^R^ backbone, using primers indicated in Table S2. In the case of the *pepB* HaloTag reporter, a version containing the coding sequence for the N-terminal regulatory domain was first constructed, and then this coding sequence was removed by using the indicated primers to amplify the plasmid lacking this sequence and recircularising the plasmid by blunt end ligation with polynucleotide kinase and T4 DNA ligase (New England Biolabs Cat # M0201S and M0202S respectively).

Reactions were transformed into *E. coli* DH5α cells by electroporation and all constructs were verified by Sanger or long read Oxford nanopore (Plasmidsaurus) sequencing. The constructs were subsequently introduced into the *attB* site in the chromosome of *P. aeruginosa* UCBPP-PA14 wild type or mutant strains by tetra-parental conjugation using strains HB101/pRK2012 and SM10/pTNS1 as a helper strains, as described above. Primers PGlmS-up and PGlmS-down were used to confirm chromosomal integration.

### SutA antibody production

A polyclonal antibody against SutA was raised by first constructing an expression plasmid encoding a GST-tagged fragment containing amino acids 46-101 of SutA. For this, two PCRs were performed using Q5 2x Master Mix (New England Biolabs, Cat # M0492S), one using primers GST_SutA46-101_plasmidF and GST_SutA46-101_plasmidR with purified pGEXP-Dendra plasmid as template to amplify the plasmid backbone, and another PCR with primers GST_SutA46-101_sutAF and GST_SutA46-101_sutAR with genomic DNA of strain PA14 WT to amplify the partial *sutA*-sequence. The resulting PCR products were ligated by Gibson assembly using the NEBuilder HiFi Assembly Mix (New England Biolabs Cat # E2621S). Ligations were transformed into *E. coli* DH5α cells by electroporation and verified by Sanger sequencing. The confirmed expression plasmid was introduced into *E. coli* strain BL21 and recombinant protein was expressed in 4x 1 L cultures that were inoculated with 12 ml of an overnight culture, and induction with 1 mM IPTG after 2 hrs growth at 30°C. Four hours after induction, 1 L cultures were pelleted by centrifugation, and each pellet was resuspended in 16 ml GST lysis buffer (50 mM Tris-HCl (pH8.0), 150 mM NaCl, 0.1 mM EDTA, plus one cOmplete protease inhibitor tablet (Sigma Aldrich) per 10 ml). Cell suspensions were sonicated at six 15 s pulses at 50% amplitude, after which the lysate was cleared by centrifugation at 40,000g, 4°C, for 45 minutes and supernatants were combined. For purification of GST- SutA46-101, 3 ml of pre-washed (2x washes in GST lysis buffer) glutathione agarose slurry (abcam) was added to the cleared lysate and the sample was incubated at room temperature for 30 mins before loading the sample into a reusable column (BioRad). Accumulated slurry was washed three times with 5 ml GST lysis buffer to remove weakly interacting proteins prior to adding 3 ml freshly made elution buffer (50 mM Tris-HCl (pH8.0), 150 mM NaCl, 0.1 mM EDTA, 10 mM Reduced (Free) glutathione) to the column. One column volume (∼750 µl) eluate was allowed to flow through, after which the column was incubated for 10 mins at room temperature. The remaining elution buffer was collected as a fraction, and the elution procedure was repeated 3-5 times. Eluted fractions were run on a gel to determine which contain protein of interest at highest quantities and purities (see Fig. S8A). 1 mg GST-SutA in elution buffer was used for antibody production by immunising two rabbits with the GST-SutA protein and collecting serum from both animals after 63 days (David’s Biotechnologie GmbH (Germany)). Sera were affinity purified against untagged SutA aa41-105 protein (SutA ΔN, see ref [15]) that was fixed to a resin. Recovered antibody was diluted 1:1 with glycerol for storage at −20 C.

### Immunoblot analysis

Bacteria were harvested at appropriate time points by centrifugation and lysed in 1× SDS loading buffer (62.5 mM Tris-HCl, pH 6.8, 5 % (v/v) glycerol, 2 % (w/v) SDS, 0.05 % (w/v) bromophenol blue, 0.25 % (v/v) β-mercaptoethanol), in a volume of OD_600nm_ value / 6.4 per ml culture. Proteins were size-separated by loading 12 µl of each lysate onto pre-cast, stain-free, 4–20% gradient gels (Mini-PROTEAN® TGX, BioRad, Cat # 4568096) and running the gels in 1x Tris-Glycine-SDS running buffer (25 mM Tris, 192 mM glycine, 0.1 % (w/v) SDS) for 65 mins at 120 V. Images of the total protein (TP) profiles of each sample were obtained using stain-free technology on a VWR Imager CHEMI Premium (Avantor) imaging system after 1 min incubation under UV light. Proteins were transferred onto nitrocellulose membranes (Amersham Protean 0.1 µm pore size, Fisher Scientific UK, Cat # 15239794) using a Mini Trans-Blot cell (BioRad, Cat # 1703930) wet transfer apparatus at either constant 100 V for 1 hr for SutA blots, or constant 350 mAmp for 1 hr for all other blots. Membranes were blocked in blocking buffer (1x TBS, 0.1 % (v/v) Tween-20, 5 % (w/v) milk) overnight at 4°C. Membranes were incubated with primary antibody in blocking buffer for 1 hr at room temperature, followed by 3× 10 min washes in TBS-T (1x TBS, 0.1 % (v/v) Tween-20), after which membranes were incubated with secondary antibody in TBS-T for 1 hr at room temperature, followed by 3× 10 min washes in TBS-T. The membranes were then incubated with chemiluminescent substrate (ECL Clarity, BioRad, Cat # 170-5061) for 5 mins at room temperature prior to imaging on an AZURE 600 scanner (Azure Biosystems). Protein gel images and immunoblot images were prepared for publication using Adobe Photoshop using the auto Tone and auto Contrast function. Specific primary antibodies included: anti-SutA (polyclonal, rabbit, used at 1:500, see production method) and anti-GFP (monoclonal, rabbit, used at 1:3,000, Abcam #ab183734). Secondary antibodies included anti-rabbit IgG (H+L) – HRP conjugate (Invitrogen, polyclonal, goat, used at 1:5,000, ThermoFisher, Cat # 31460).

Signal intensities were measured using the Gel Analysis function of the Fiji_ImageJ image analysis software (Version 1.54f [20]). All lanes were internally normalised to an in-gel reference sample from the signal intensities of the entire lane of the corresponding Total Protein gel image. All GFP or SutA signal intensities were normalised across different blots by including the same reference sample (PA14 WT grown in LB to stationary phase for SutA blots or WT plus pSG-P reporter grown in LB overnight for GFP blots) on each gel.

### BONCAT Proteomics

BONCAT proteomics was performed as previously described[7] with minor modifications. Briefly, WT and *ΔsutA* cells carrying the NLL-metRS mutant tRNA synthetase under control of the P_trc_ promoter, integrated into the *attB* site on the chromosome[66], were grown into stationary phase overnight, then diluted to OD_500_=0.005 in 35 mL LB and grown shaking in a flask at 37 C for 24 hours into stationary phase again. Cells were collected by centrifugation then washed and resuspended at approximately OD_500_=4 in phosphate-buffered minimal media with no carbon or nitrogen source. This was intended to maintain the high cell density but mitigate changes to pH that typically occur in late stationary phase. After 16 hours, ANL was added to a concentration of 500 µM. Following labelling for 24 hours, 200 µg/ml chloramphenicol was added to cultures to terminate protein synthesis before pelleting, washing in PBS and flash freezing. Pellets were lysed in 2% SDS, 100 mM Tris pH 8.0, 100 mM iodoacetamide, 0.2 μL/mL benzonase with cOmplete EDTA-free protease inhibitor (Sigma Aldrich); clarified by spinning at high speed in a benchtop centrifuge (13,000 g for 10 minutes); and normalised to equivalent protein concentrations. Urea buffer (8M Urea, 150 mM NaCl, 1X protease inhibitor) and 25 μL of DBCO agarose beads (Jena Bioscience) were added to normalised samples and rotated for four hours at room temperature. Beads were collected by centrifugation at 1000 rpm, washed with SDS wash buffer (0.8% SDS, 150 mM NaCl, 100 mM Tris pH 8.0) and resuspended in SDS wash buffer with 5 mM dithiothreitol at room temperature for 30 min. Beads were then incubated in SDS wash buffer with 100 mM iodoacetamide at 50°C in the dark for 45 minutes. Beads were washed in SDS wash buffer and transferred to Poly-Prep chromatography columns (BIO-RAD). Columns were washed 8 times with 5 mL SDS wash buffer, 8 times with 5 mL urea buffer with 100 mM Tris pH 8.0, and 8 times with 5 mL 20% acetonitrile. Finally, beads were resuspended in 10% acetonitrile with 50 mM ammonium bicarbonate and submitted to the University of Dundee Fingerprints Proteomics Facility for analysis. Peptides were recovered from the beads and supernatant of the submitted samples by tryptic digest overnight at 37°C with 0.75 µg of trypsin per sample. The digest was repeated for 6 hours the following day. Digested peptides were resuspended in 1% formic acid (FA) and an equivalent of 1µg was run on a Q-Exactive Plus instrument *(Thermo Scientific)* coupled to a Dionex Ultimate 3000 HPLC system *(Thermo Scientific)* with LC buffers comprising of buffer A (0.1% FA) and buffer B (80% ACN, 0.1% FA). Peptides from each sample were loaded at 10 μL/min onto a trap column (100 μm × 2 cm, PepMap nanoViper C18 column, 5 μm, 100 Å, Thermo Scientific) equilibrated in 0.1% TFA for 15min. The trapping column was washed for 6 min at the same flow rate with 0.1% TFA and then then switched in-line with a Pharma Fluidics, 200 cm, µPAC nanoLC C18 column, equilibrated with buffer A at a flow rate of 300nl/min for 30 min. The peptides were eluted from the column at a constant flow rate of 300 nl/min with a linear gradient from 1% buffer B to 3.8% buffer B in 6 min, from 3.8% B to 12.5% buffer B in 40 min, from 12.5% buffer B to 41.3% buffer B within 176 min and then from 41.3 % buffer B to 61.3% buffer B in 34 min. The gradient is finally increased from 61.3% buffer B to 100% buffer B in 10 min, and the column was then washed at 100% buffer B for 10 min. Two blanks were run between each sample to reduce carry-over. The column was kept at a constant temperature of 50°C.

Raw data was acquired in positive and Data Independent Acquisition (DIA) mode using an easy spray source. The source voltage was set to 2.2 Kv and the capillary temperature was at 275°C. Scan cycle comprised a full MS scan with an m/z range of 345-1155, resolution of 70,000, Automatic Gain Control (AGC) target 3×10^6^ and a maximum injection time of 200ms. MS scans were followed by DIA scans of dynamic window widths. DIA spectra were recorded with a first fixed mass of 200m/z, resolution of 17,500, AGC target 3×10^6^ and a maximum IT of 55ms. Normalised collision energy was set to 25 % with a default charge state set at 3. Data for both MS scan and MS/MS DIA scan events were acquired in profile mode. Thermo RAW files were analysed using Spectronaut version 17 *(Biognosys)* using directDIA. Samples were searched against the UCBPP-PA14 proteome (Uniprot reference 20220523) with variable modifications of Oxidation(M), Dioxidation(MW), Acetyl (Protein N-term), Deamidation (NQ) and Gln->Pyro-Glu set and a fixed modification of Carbamidomethyl (C). For identification a precursor PEP (Posterior Error Probability) cutoff of 10%, a protein Qvalue cutoff of 5%, and a false discovery rate of 1% were used. Quantification was done using the protein LFQ method Quant 2.0 (SN standard) within Spectronaut. Output is provided in Table S1. To construct Fig. 5B, the RpoS and FliA regulon lists (locus tags) from Schulz *et al*. were searched against the identified proteins in the proteomic data set. Uniprot accession numbers associated with each relevant locus tag were extracted from the supplemental file of Munro *et al.*[7] For every gene whose protein was robustly identified in our proteomics data set, we also extracted RNA-Seq and ChIP-Seq data from Supplemental table S3 of Babin *et al.*[9].

### Microscopy

All microscopy was performed using live bacteria spotted onto agarose pads (1.2 (w/v)% agarose in 1x PBS) within a Gene Frame (ThermoFisher, Cat # AB0577) as described in ref[7]. Briefly, phase contrast and fluorescent images were acquired using a Nikon Eclipse Ti2 microscope fitted with a 60X phase contrast objective. Fluorescence of interest was recorded with the following settings: GFP (excitation 470 nm LED, emission 510 FITC nm filter as part of a Pinkel Quad filter cube) or Janelia Fluor 646 (JF646; excitation 640 nm LED, emission 700 nm Cy5 filter as part of a Pinkel Quad filter cube). Phase-contrast images used a 20-100 msec exposure time, while JF646 images were acquired using 600 or 700 msec exposure time. GFP fluorescence settings were calibrated to the brightest sample within each experiment but were usually 60-100 msec exposure time. Intensities were normalised to a reference sample collected on the same day. All microscopy images were processed in ImageJ, using the plugin MicrobeJ to first identify and segregate individual bacteria from the phase contrast image and to then quantify average cellular fluorescence intensities. At least three fields of view per sample containing 200-1,200 individual bacterial cells were analysed.

### RNA extractions

RNA was extracted from bacterial cultures at appropriate time points after pelleting 1.5 ml of exponential growth stage samples (OD_600nm_ < 1), 1 ml of transition-phase samples (OD_600nm_ 1-3), or 0.5 ml of stationary growth stage samples (OD_600nm_ > 3), and snap freezing pellets in liquid nitrogen. Pellets were stored at −80°C until RNA extraction using the Monarch® Total RNA Miniprep Kit (New England Biolabs, Cat # T2010L) according to the manufacturer’s recommendations. In brief, pellets were resuspended in 200 µl lysozyme buffer (2 mg/mL lysozyme in 10 mM Tris-HCl, 1 mM EDTA, pH 7.0 (Thermo Fisher, Cat # AM9861)) and incubated for 5 mins at room temperature. Cells were lysed by adding 2 volumes RNA Lysis Buffer and the lysate was passed through gDNA removal columns and RNA binding columns as described in the product manual. Total RNA was eluted in 50 µl nuclease-free water.

Contaminating gDNA was removed using the Invitrogen TURBO DNA-free*™* Kit (Thermo Fisher. Cat # AM1907), following the rigorous treatment protocol. DNase enzyme and salts were removed using the Monarch^®^ Spin RNA Cleanup Kit (500 μg, New England Biolabs, Cat # T2050L). DNA-free RNA was eluted in 50 µl nuclease-free water. Samples were verified to be free of genomic DNA by standard PCR, with genomic DNA as a positive control.

### Reverse transcription and qPCR

0.5-1 μg of DNA-free RNA was reverse transcribed to generate cDNA using random hexamer primers (Promega, Cat #) or gene-specific primers containing a tag-sequence overhang for single-stranded RT-PCR and Invitrogen Superscript III Reverse Transcriptase Synthesis System (Thermo Fisher, Cat # 18080093) according to the manufacturer’s recommendations, including an extended synthesis step for 60 mins at 50°C, followed by RNaseH digest at 37°C for 30 min. qPCR was performed using the GoTaq® qPCR (Promega, Cat # A6001) 2x SYBR™ Green premix and a primer complementary to the tag sequence included in the cDNA and a matching forward (fw) primer annealing to internal regions of the target genes (Supplementary Table S2). Reactions (40 cycles with an annealing temperature of 60 °C) were analysed on a StepOnePlus Real-Time PCR System (Thermo Fisher) using a dilution series of *P. aeruginosa* PA14 wt gDNA as a standard curve and the *oprI* gene as a reference gene for normalisation. Measurements were made on two biological replicates with three technical replicates each.

### Primer walking

Primer walking was performed on single-stranded cDNA molecules generated as described above. Standard PCR reactions using the GoTaq® G2 DNA Polymerase (Promega, Cat # M7845) enzyme and buffers using the same tag primer combined with a single fw primer per reaction. Primer binding sites within the internal and promoter regions of the target genes were spaced 20-40-bp apart. PA14 wildtype gDNA was used in control PCR reactions using the same fw primers but combining it with the original gene-specific RT-primer that was used to create the cDNA molecules instead of the tag primer. PCR products were separated on 2% (w/v) agarose gels in TAE buffer and visualised using Invitrogen SYBR™ Safe DNA Gel Stain (Thermo Fisher, Cat # S33102) and a VWR Imager CHEMI Premium (Avantor) imaging system.

### 5’ RACE

Transcription start site mapping was performed on single-stranded cDNA molecules generated as described above and using terminal transferase enzyme, as described in the supplementary information for Bergkessel *et al.* [15]. In brief, in 10 µg of DNase-treated RNA from PA14 wildtype grown to exponential phase in LB medium was used for reverse transcription using 4 pmol *sutA*-specific RT primer, 500 μM dNTPs, 5 μM DTT, 1x reverse transcriptase buffer, and 300 units SuperScript III reverse transcriptase in a 40 μl reaction. Primer binding was allowed to occur for 5 min at 65°C prior to addition of reverse transcriptase mixture and the reaction allowed to proceed for 45 min at 55 °C. The reaction was stopped by incubation for 15 min at 70°C, after which 2 units RNaseH enzyme were added, and the reactions were incubated at 37°C for 20 min to degrade RNA-DNA hybrids. The cDNA was cleaned up using the Monarch DNA clean-up kit (New England Biolabs, Cat # T1030S) and eluted in 20 µl nuclease-free water. Poly-C tails were added to the 3’ end of the cDNA using terminal transferase (TdT, New England Biolabs, Cat # M0315S) in 50 µl reactions containing 18 µl cDNA, 1x TdT buffer, 0.25 mM CoCl_2_, and 5 µl of a mix containing 9.5 mM dCTP and 0.5 mM ddCTP. Reactions were incubated at 37°C for 1 hr and heat inactivated for 10 mins at 70°C. Unused CTPs were removed by passing the C-tailed cDNA through a Monarch DNA clean-up kit (New England Biolabs) column and eluting in 20 µl nuclease-free water. The resulting C-tailed cDNA was then used as a template in a first round PCR reaction using GoTaq G2 polymerase (Promega) with a target-specific primer (RACE_sutA_PCR1-rv) and a C-tail-specific primer (RACE-PCR1-fw), both of which add on additional overhang sequences. This PCR product was then used as the template in a second round PCR reaction with primers complementary to the added overhang sequences that were part of the primer in the first round PCR (RACE-PCR2-fw&rv) and 1 µl of the product of the first PCR as template. The products of the second PCR round were separated on 2% (w/v) agarose gels in TAE buffer and visualised using Invitrogen SYBR™ Safe DNA Gel Stain (Thermo Fisher, Cat # S33102) and a VWR Imager CHEMI Premium (Avantor) imaging system. Individual PCR fragments were excised from the gel, dissolved in Monarch® Buffer BY (New England Biolabs, Cat # T1121L), purified using the Monarch DNA clean-up kit (New England Biolabs), and eluted in 12 µl nuclease-free water. PCR products were then cloned into a pGEM®-T Easy using the TA cloning vector system (Promega, Cat # A1360) using 50 ng plasmid and a 3:1 molar ratio vector to insert, depending on the estimated size of each product and according to the manufacturer’s recommendations. Ligated plasmids were transformed into CaCl_2_-competent *E. coli* DH5α cells and insert-containing plasmids were selected for by blue-white screening on LB plates supplemented with X-Gal, IPTG and ampicillin. Plasmids from individual clones were extracted using the Monarch® Spin Plasmid Miniprep Kit (New England Biolabs, Cat # T1110) and inserts were Sanger sequenced using primer AS1-fw as sequencing primer.

### Northern blotting

Non-radioactive northern blotting was performed using the Invitrogen NorthernMax™-Gly Kit (Thermo Fisher, Cat # AM1946) and Biotin-modified RNA probes. Short (100-120-bp), complementary regions within each target gene were PCR amplified using Q5 2x Master Mix (New England Biolabs) and primer pairs listed in Table S2, one of which included an overhang containing a T7 class III phi6.5 promoter. Randomly Biotin-modified RNA probes were created using Biotin-16-UTP and T7-mediated *in vitro* transcription using 0.5 µg of PCR as template DNA and the HighYield T7 Biotin16 RNA Labeling Kit (Jena Bioscience, Cat # RNT-101-BIO16) according to the manufacturer’s recommendations. Template DNA was removed using the Invitrogen TURBO DNA-free*™* Kit (Thermo Fisher. Cat # AM1907), following the rigorous treatment protocol. DNase enzyme and salts were removed using the Monarch^®^ Spin RNA Cleanup Kit (500 μg, New England Biolabs, Cat # T2050L). RNA probes were eluted in 50 µl nuclease-free water and stored at −80°C until used.

For the northern blot, 30 μg of DNase-treated total RNA was denatured by heating in glyoxal/DMSO Loading Solution and then separated alongside lanes containing either Millennium™ RNA Marker (Thermo Fisher Cat # AM7150) or RiboRuler Low Range RNA Ladder (Thermo Fisher, Cat # SM1831) on a 1.5 % agarose-LE gel prior to blotting onto a BrightStar™-Plus Positively Charged Nylon Membrane by capillary transfer, following the manufacturer’s recommendations. RNA was crosslinked to the membrane using a UV crosslinker CL-508 (Uvitec Cambridge) at a dosage of 0.120 J/cm². The membranes were pre-hybridised in ULTRAhyb™ Hybridization Buffer for >30 min at 68°C, after which individual RNA probes were added at a concentration of 25 nM. Membranes were hybridised with the probes at 68°C overnight. Washes in Low Stringency and High Stringency Wash Solution were performed according to the kit protocol. Membranes were subsequently blocked in Intercept® (PBS) Blocking Buffer (Licor, Cat # 927-70001) supplemented with 1% (w/v) SDS for 3 hours, after which fresh blocking buffer containing 1:5,000 diluted Streptavidin- IRDye 800CW conjugate (Licor, Cat # 926-32230) was added to the membranes and incubated in the dark for 45 min. Membranes were washed three times in 1X PBS-T (0.1% (v/v) Tween-20) for 5 minutes each, followed by a wash in 1X PBS for 5 minutes at room temperature. Membranes were imaged on an AZURE 600 scanner (Azure Biosystems) using fluorescent IRDye 800CW settings. Where necessary, membranes were stripped by submerging them in a bottle containing 0.1% (w/v) SDS in DEPC-treated water and autoclaving in a benchtop autoclave on a standard cycle (120°C, 15 min).

Signal intensities were measured using the Gel Analysis function of the Fiji_ImageJ image analysis software (Version 1.54f [20]). All lanes were normalised using signals from the same membrane after rehybridization with an *oprI* reference probe.

### FliA purification

FliA was purified as previously described[41] with some modifications. E. coli BL21(DE3) were transformed with the pET15b vector carrying 6His-TEV-FliA. Cells were grown in 800 ml Terrific Broth plus 100 ug/ml carbenicillin in a 2L flask shaking at 37 C until OD_600_=0.5 and then IPTG was added at 0.5 mM to induce expression at 20 C for 8 hr. Cells were collected by centrifugation and frozen at −70 C. They were then resuspended in 20 ml cold lysis buffer (50 mM Tris HCl pH 8.0, 300 mM NaCl, 10 mM imidazole, 5% glycerol, complete mini protease tablet (Roche)) and lysed by ultrasonication, and lysate clarified by spinning at 20,000 rpm for 45 minutes in a JA25.5 rotor. Clarified lysate was loaded onto a 5 ml HisTrap column (GE Healthcare), washed with a further 20 ml lysis buffer, then eluted with a gradient to elution buffer (50 mM Tris HCl pH 8.0, 300 mM NaCl, 400 mM imidazole, 5% glycerol). Fractions 7-11 were pooled and buffer exchanged by centrifugal filtration (Amicon) into TEV cleavage buffer (50 mM Tris, 0.5 mM EDTA, 1 mM DTT)(Fig. S8B). Protein concentration was measured by BCA assay and TEV protease was added at a mass ratio of 1:50. Cleavage was allowed to proceed overnight at 4 C. Protein was then dialysed into storage buffer (20 mM Tris HCl pH 8.0, 150 mM NaCl) and glycerol was added to a final concentration of 20%. Aliquots were snap frozen in liquid nitrogen and stored at −70 C.

### *In vitro* transcription assays - fluorescence

A fluorescent *in vitro* transcription assay using mango array technology was performed based on the method described by He *at al*. [16], with some modifications. DNA templates were produced by PCR using primers Tn7-chk-fw & Tn7-chk-rv (*fliC*-mango and *pepB*-mango templates) or PW-8-fw & Tn7-chk-rv (*sutA-*mango template) in standard PCR reactions using the GoTaq® G2 DNA Polymerase (Promega, Cat # M7845) kit and 1 µl per 50 µl PCR reaction template DNA consisting of purified plasmid DNA of pUC18-Tn7 constructs encoding the respective promoter / gene fusions to a mango II array. The resulting PCR products were 1361-bp (P*_fliC_*::5x-mango), 1550-bp (P*_pepB_*::6x-mango) or 1629-bp (P*_sutA_*-prox-*sutA*::6x-mango) in length.

Initial tests revealed high background fluorescence in samples that contained templates and buffer, but where no RNA polymerase was added, which we partially attributed to single-stranded DNA being present in the PCR products. Therefore, 8 µl of thermolabile exonuclease I (New England Biolabs, Cat # M0568S) was added directly to 50 µl PCR products in order to remove ssDNA, and reactions were incubated at 37°C for 4 mins, followed by heat inactivation at 80°C for 1 min.

*P. aeruginosa* RNAP holoenzymes were created by mixing purified RNAP core complex with either purified RpoD (1:3 molar ratio), RpoS (1:5 molar ratio), or FliA (1:5 molar ratio) and incubating for 15 mins at 37°C (see reference [15] for details on purification of RNAP, RpoD ( σ^70^), and RpoS (σ ^S^), and this methods section for purification of FliA). See Fig. S7C for a gel image of all purified proteins used in this assay.

*In vitro* reactions were performed in volumes of 20 μl and contained RNAP holoenzymes (final concentration: 40 nM), *exo*I-digested template DNA (final concentration: 30 nM), purified SutA (final concentration: 0 nM or 125 nM; see ref [15] for details on purification of SutA protein), in modified 1x reaction buffer (50 mM Tris-acetate pH 8.0, 100 mM K-acetate, 10 mM Mg-acetate, 1 mM DTT, 5% (v/v) glycerol, and 0.01% (v/v) Tween-20) in 384-well plates with flat, transparent bottoms and black walls (Greiner, Cat # 781096). Reactions were incubated for 10 mins at 37°C to allow open complex to form, after which an NTP mixture was added at a final concentration of 0.2 mM of each nucleotide. Plates were incubated for 30 mins at 37°C to allow for transcription to occur before addition of TO1-3PEG-Desthiobiotin fluorophore (Caltag Medsystems, Cat # ABM-G7956) at a final concentration of 0.5 µM. Plates were incubated at 37°C inside a Tecan Spark platereader, shaken for 60 sec at low amplitude to mix the samples, and fluorescence intensities were collected every 3 mins at an excitation wavelength of 485 nm and an emission wavelength of 535 nm. Data acquisition was stopped when fluorescence reading remained steady. In all assays, a no-RNAP control sample was included, and these background fluorescence values were subtracted from *in vitro* transcription reaction fluorescence values. The assay was performed on three different days, with 2-3 technical replicates per holoenzyme – template combination, which were averaged to give one value per day and these values are plotted.

### *In vitro* transcription assays - radioactivity

We produced linear templates of 120-170 bp containing the *pepB* promoter and 42-50 bp of transcript sequence for use in single-turnover initiation experiments with or without SutA. dsDNA templates were prepared by PCR from plasmids carrying the relevant promoter sequences or directly from *Pseudomonas aeruginosa* UBCPP-PA14 genomic DNA, using the Kappa high-fidelity hot-start 2x master mix according to the manufacturer’s instructions (see strain and primer tables for plasmid and primer details). PCR products were checked by electrophoresis on 2% agarose gels to ensure that they consisted of a single product, purified from primers and residual dNTPs using the DNA Clean and Concentrator kit (Zymo Research), and quantified by NanoDrop (Thermo Fisher). Reactions were assembled as follows: RNAP holoenzyme (40 nM final concentration), DNA template (15 nM final concentration), TGA buffer (20 mM Tris-acetate pH 8.0, 2 mM Na-acetate, 2 mM Mg-acetate, 4% glycerol, 0.1 mM DTT, 0.1 mM EDTA), and water were mixed in a volume of 3 μl and added to 1 μl SutA (at 5x the final concentration) or storage buffer on ice. These 4 μl reactions were incubated for 16 min to allow open complex to form. 1 μl NTP mix (375 μM initiating dinucleotide (ATP), 250 μM each NTP not carrying 32P label (UTP and GTP), 100 μM cold NTP of the same type as that carrying the label (CTP), 0.75 μCi α^32^P CTP (3000 Ci/mmol, 10 mCi/ml, Perkin Elmer), and 100 μg/ml heparin) was added and transcription was allowed to continue for 8 minutes before reactions were quenched. Reactions were quenched with an equal volume of urea stop buffer (8 M urea, 10 mM EDTA, 0.8x TBE, 2 mg/ml bromophenol blue, 2mg/ml xylene cyanol FF, 2 mg/ml amaranth) and heated to 95 °C for 2 min immediately before gel loading. 20% acrylamide denaturing Urea-TBE gels were prepared using the Sequa-gel system (National Diagnostics) according to the manufacturer’s instructions except TBE was added to 0.5x instead of 1x. A 60-well comb was used and gels were run using the Owl S3 vertical sequencing gel system (Thermo Fisher). 2 μl sample was loaded per lane. After electrophoresis, one glass plate was removed and the gel was covered with plastic wrap and exposed directly to the phosphorimager screen (Molecular Dynamics) for 12-48 hr. Screens were imaged with a Typhoon FLA 9000 gel imaging system and images were analysed using the ImageJ gel lane tool[20].

### rRNA FISH

Per cell ribosome amounts were measured by hybridisation of either a Cy5-labeled EUB338 oligonucleotide probe against the 16S rRNA or a control “scramble” probe (Integrated DNA Technologies), against the rRNA in fixed cells, as previously described[7]. Briefly, cells were grown into stationary phase overnight, then diluted to OD_500_=0.005 and allowed to grow for 24 hours into stationary phase again. Cells were collected by centrifugation then washed and resuspended at approximately OD_500_=4 in phosphate-buffered minimal media with no carbon or nitrogen source. This was intended to maintain the high cell density but mitigate changes to pH that typically occur in late stationary phase. ANL was added for consistency with the BONCAT experiment described above. After 32 hrs incubation, cells were fixed in 4% paraformaldehyde in PBS for 30 minutes, permeabilized with ice cold ethanol and washed three times in 0.9% NaCl. They were then hybridized for 3 hours at 46°C in hybridization buffer (900 mM NaCl, 20 mM Tris pH 7.6, 0.01% SDS, 20% HiDi formamide and 2 μM fluorescent probe). Samples were then washed twice in wash buffer (215 mM NaCl, 20 mM Tris pH 7.6, 5 mM EDTA) for 15 minutes at 48°C shaking in the dark and resuspended in PBS plus 10 ug/ml DAPI for use in flow cytometry. Samples were analysed on an LSRFortessa flow cytometer (BD Biosciences) and 50,000 events were collected for each sample. DAPI fluorescence was measured using 355 nm excitation and emission was collected at 450/50 nm. Cy5 fluorescence was detected using 640 nm excitation and emission was collected at 670/14. Cytometry data was imported and analysed in FlowJo V10.7.2 (Becton Dickinson). Populations were first gated based on forward-scatter and DAPI profile to identify DNA-containing single cells. The pulse height in the Cy5 channel was then collected for the gated subpopulation, which averaged 50-70% of all events. For each replicate, the average scramble probe intensity was subtracted from the average EUB338 probe intensity to plot mean background-subtracted values in bar charts. Histograms for all EUB338 probe replicates and a representative scramble probe replicate are shown.

### Halo tag assays

HaloTag reporters were used to measure per-cell new protein synthesis during stationary phase. Strains expressing the halo-tagged target proteins as well as untagged control strains (PA14 WT and PA14 *ΔsutA*) were grown in triplicates in 50 ml LB medium for 14 hrs at 37°C with aeration before a dark ligand (50 µM 6-chloro-hexanol in LB medium) was added to each culture to quench pre-existing reporter proteins and cultures were incubated for a further 30 mins at 37°C in the shaking incubator. Unquenched samples were taken from one replicate each prior to addition of the dark ligand. In the meantime, Janelia Fluor® 646 Halo Ligand (Promega, Cat # HT106A) was dissolved in filtered, spent culture supernatant from either PA14 WT or PA14 *ΔsutA* (prior to the addition of dark ligand) to give a 1x final concentration of 200 nM. 200-300 µl of each quenched culture was transferred to microfuge tube and cells were washed twice in 1x PBS before pelleting. Cell pellets were resuspended in 0.83x sample volumes in 1x fluorescent Halo ligand and transferred to 7 ml Bijou tubes, which were incubated at 37°C with aeration. At each designated time point (T0, T4, T8), 50 µl samples were taken from each Bijou, and cells were washed once in 1 ml 1x PBS before resuspending the pellets in 500 µl 1x PBS. 1 µl of each sample was used directly for microscopy as described. Images were analysed suing the MicrobeJ plug-in after background subtraction in the fluorescent channel using a rolling ball radius of 50 pixels. Fluorescence intensity density plots were created using the geom(density) aesthetic in ggplot2[67], whereas distributions were fitted using the fitdistrplus library[67] in Rstudio (2025.05.1 Build 513 © 2009-2025 Posit Software, PBC, “Mariposa Orchid” Release (ab7c1bc7, 2025-06-01) for windows). The original, unfiltered dataset was used to calculate median florescence values per sample as well as for the density plots, whereas a filtered dataset where extreme outliers were removed (using GraphPad Prism ‘remove outliers’ function by ROUT method with Q = 0.1 %; this removed 11,333 datapoints (3.9 %) from a dataset containing 289,778 individual values) was used for fitting of distributions.

### Swimming Assay

Flagellar motility was measured as previously described with minor modifications[7]. Triplicate overnight starter cultures of WT and *ΔsutA* bacterial were diluted to OD_500_ = 0.005 and grown for 15 hours with shaking in universal tubes at 37° C. 50 µl of culture was then placed into a chambered cover slip (ibidi, Cat # 81816) while maintaining the temperature at 37°, and then the chambered cover slip was rapidly moved into a heated microscope enclosure. Focusing at the surface of the coverslip using a 60x oil phase-contrast objective of a Nikon Eclipse Ti2 microscope, such that the most coverslip-proximal bacteria were phase bright, films lasting 30 seconds and containing 91 frames (3 fps) were generated. The ImageJ plugin TrackMate was used to analyse phase contrast films by identifying individual cells and tracking their displacement between frames of the film [68]. Briefly, the “thresholding” detection method was used to identify cells in each frame. Identified cells were filtered using an automatic “quality” threshold. Tracks of individual cells across the frames were calculated using the default “Simple LAP Tracker” with “Linking max distance” and “gap-closing max distance” both set to 15 microns, and “gap-closing max frame gap” set to 2. For each film, tracks lasting less than 2 seconds were discarded. The experiment was performed twice, and all tracks from the 6 total replicates performed lasting longer than 2 seconds were aggregated. The population density for mean speed over the track length per cell was plotted for both strains using ggplot2 “geom_density” in Rstudio.

### Milk casein halo assay

To prepare milk plates, 7.5 g agar was added to 400 ml dH_2_O and 7.5 g non-fat dry milk powder was added to 100 ml dH_2_O. The agar was autoclaved (20 minutes at 121 C) while the milk was pasteurised (20 minutes at 65 C), then the two solutions were mixed and plates were poured to achieve final concentrations of 1.5% w/v agar and milk.

Strains were streaked from freezer stocks and overnight cultures were started in triplicate from distinct colonies. Stationary phase cultures were normalised to equivalent optical densities by spinning aliquots of culture and removing supernatant from cultures with lower OD measurements. 4 µl from each normalised culture was spotted onto milk plates and dried. Plates were incubated at 37° C for 18 hours and then photographed.

### Promoter sequence analysis

RpoD and transcription factor binding sites were predicted using SAPPHIRE [26] or Prodoric [30], respectively. The PRODORIC Virtual footprint function was used, and matrices for *P. aeruginosa* RhlR (MX000011), LasR (MX000012), IHF (MX000010), MexR (MX000001), Anr (MX000002), FleQ (MX000104), AlgR (MX000101), GlpR (MX000105), OxyR (MX000107) and PhhR (MX000108) and E. coli Lrp (MX000096) were searched against the upstream sequence included in the pSG-P reporter construct. Matches to the coding strand were reported. In the case of Lrp, results were filtered to scores above 1.65, which corresponded to 0.72 of the maximum score.

For assessing conservation of the regulatory elements, the SutA protein sequence was submitted to the GeCoViz[69] website to assess the genomic neighborhoods of other members of the same eggnog orthologous group (33HP8). Representative genomes across the Pseudomonadaceae family were chosen for display in Fig. S4. To investigate in more detail the specific sequence elements, the intergenic regions between the *amtB* and *sutA* genes were retrieved from the pseudomonas.com website for seven species. Sequences were aligned using Clustal OMEGA[70], which captured the conservation of the RpoD consensus near the beginning of these sequences. The alignments were then manually adjusted to focus on the other regulatory elements of interest, accommodating the variable lengths of the intergenic regions lacking the *yjbQ* ORF. Finally, each of the sequences was searched using the Lrp matrix (MX000096) in the PRODORIC virtual footprint function[30], and the same score cutoff was applied as for the original PA14 analysis.

### Statistics and Figures

All statistical tests were performed in GraphPad Prism. Adjusted p values are reported as ns = P > 0.05, * = P ≤ 0.05, ** = P ≤ 0.01, *** = P ≤ 0.001, ****= P ≤ 0.0001. Figures were constructed using GraphPad Prism or Rstudio with the ggplot2 package[71].

## Acknowledgments

We are grateful for the Δ*rpoS* and NLL*metRS*-expressing strains gifted to us by Dianne Newman, and for her support at the beginning of this project. We appreciate assistance from the University of Dundee Flow Cytometry and Cell Sorting Facility, Imaging Facility, and Fingerprints Proteomics Facility. We would like to thank Daniel Neill, Sarah Coulthurst, Findlay Munro, and Lisa Racki for helpful feedback and discussions. We would also like to thank Peter McBride for reagents used during the purification of FliA.

## Supporting information captions

### Supplemental Figures

**Figure S1. Culture conditions and investigation of SutA cleavage**

**Figure S2. *sutA* transcriptional units**

**Figure S3. GFP expression in modified *sutA-gfp* reporter strains**

**Figure S4. *sutA* genomic context and regulatory motifs are conserved across Pseudomonadaceae**

**Figure S5. Representative SutA western blots and statistical analysis of WT and putative regulator mutants**

**Figure S6. Effects of regulatory mutants on growth, SutA levels and the production of pyocyanin**

**Figure S7. Supporting data for in vitro transcription and FISH experiments**

**Figure S8. Fractions collected during purifications of proteins newly purified for this study**

**Figure S9. Description of HaloTag fluorescence distributions and average intensities**

**Figure S10. Analysis of HaloTag distributions**

### Supplemental Tables

**Table S1. Proteomics data**

**Table S2. Strains, plasmids, and primers used in this study**

